# A Versatile Anti-CRISPR Platform for Opto- and Chemogenetic Control of CRISPR-Cas9 and Cas12 across a Wide Range of Orthologs

**DOI:** 10.1101/2024.11.25.625186

**Authors:** Luca Brenker, Sabine Aschenbrenner, Felix Bubeck, Kaloyan Staykov, Carolin Gebhardt, Benedict Wolf, Michael Jendrusch, Ann-Sophie Kröll, Jan Mathony, Dominik Niopek

## Abstract

CRISPR-Cas technologies have revolutionized life sciences by enabling programmable genome editing across diverse organisms. Achieving dynamic and precise control over CRISPR-Cas activity with exogenous triggers, such as light or chemical ligands, remains an important need. Existing tools for CRISPR-Cas control are often limited to specific Cas orthologs or selected applications, restricting their versatility. Anti-CRISPR (Acr) proteins, natural inhibitors of CRISPR-Cas systems, provide a flexible regulatory layer but are constitutively active in their native forms.

In this study, we built on our previously reported concept for optogenetic CRISPR-Cas control with engineered, light-switchable anti-CRISPR proteins and expanded it from ortholog-specific Acrs towards AcrIIA5 and AcrVA1, broad-spectrum inhibitors of CRISPR-Cas9 and -Cas12a, respectively. We then conceived and implemented a novel, chemogenetic anti-CRISPR platform based on engineered, circularly permuted ligand receptor domains of human origin, that together respond to six different, clinically-relevant drugs. The resulting toolbox achieves both optogenetic and chemogenetic control of genome editing in human cells with a wide range of CRISPR-Cas effectors, including type II-A and -C CRISPR-Cas9s, and -Cas12a. In sum, this work establishes a versatile platform for multidimensional control of CRISPR-Cas systems, with immediate applications in basic research and biotechnology and potential for therapeutic use in the future.

## INTRODUCTION

The CRISPR (clustered regularly interspaced short palindromic repeats) technologies revolutionize(d) the biosciences and hold enormous potential for the development of personalized and targeted therapies (1). At the core of this technology are CRISPR-Cas nucleases that naturally occur as part of the CRISPR adaptive immunity in prokaryotes and destroy invading nucleic acids via targeted cleavage (2). Importantly, Cas nucleases can be heterologously expressed in bacterial, plant, and mammalian cells and be programmed to target specific sequences on genomic DNA simply by providing a complementary guide RNA (3,4). Using various CRISPR-Cas9 and -Cas12 orthologs (5) from the class II, type II and -V clades, respectively, researchers can nowadays achieve highly efficient and cost-effective genomic knock-outs or knock-ins across organisms. Moreover, nuclease-compromised CRISPR-Cas variants, such as Cas nickases or catalytically inactive Cas variants (deadCas9 or deadCas12a), further broaden the application scope of CRISPR. They enable controlled expression of endogenous genes via recruitment of transcriptional activators/repressors and epigenetic modifiers (6,7), editing of DNA bases via engineered CRISPR base editors (8,9) as well as inserting and re-writing short stretches of DNA by prime editing technologies (10) .

The efficient, targeted and safe application of the CRISPR method in its various flavors demands control mechanisms to ensure the Cas enzymes are active only at the desired place, e.g. within selected cells or tissues of an organism (11,12). Moreover, for dynamic genome perturbations or to avoid unintended modifications of genomic loci which are similar in sequence to the actual target locus, it is crucial to control the activity of Cas enzymes in time (13–15).

In the past, various studies therefore focused on engineering conditional variants of CRISPR-Cas enzymes, whose activity depend on the presence of exogenous triggers such as chemicals (16–18), temperature (13) or light (13,19–21). Complementary efforts developed caged, stimulus-dependent single guide (sg)RNAs (22,23). However, all of these approaches require users to employ specific Cas or sgRNA variants and are often either (i) irreversible, (ii) restricted to a single Cas ortholog or (iii) limited to one specific application, e.g. genome editing or gene activation.

Anti-CRISPR (Acr) proteins, bacteriophage-derived, potent CRISPR-Cas inhibitors, represent a powerful, naturally occurring regulatory layer for CRISPR control. First described in 2013 by Alan Davidson’s group (24), a variety of Acrs have since been discovered that target class II, type II (25–33) and type V (34,35) CRISPR effectors as well as CRISPR-Cas13, a type VI effector (36). Remarkably, some Acrs were found to exhibit broad-spectrum activity, i.e. they can be used to effectively block the function of various different Cas9 orthologs (27,36,37).

One prominent example is AcrIIA5, a broad spectrum type II inhibitor, which targets practically all of the widely-used Cas9 orthologs such as *Streptococcus pyogenes* (*Spy*) and *Staphylococcus aureus* (*Sau*) Cas9 from the type II-A and *Neisseria meningitides* (*Nme*) Cas9 from the type II-C subtypes. AcrIIA5 is characterized by an α/β fold with an N-terminal intrinsically disordered region essential for inhibition (37). Its primary mode of action likely involves blocking Cas9 nuclease activity, though interference with DNA target binding was also observed by us (38) and others (26). AcrVA1, in turn, is a broad-spectrum inhibitor of Cas12a, acting as both, a PAM mimic as well as an RNase cleaving the Cas12a-loaded crRNA. Thereby, AcrVA1 potently blocks the activity of various Cas12a orthologs, including those from *Moraxella bovoculi* (*Mb*Cas12a), *Lachnospiraceae bacterium* (*Lb*) and *Acidaminococcus sp*. (*As*) (34,35).

Very importantly, many Acrs remain functional when heterologously expressed in eukaryotes, including human cells, therefore providing powerful building blocks for the construction of CRISPR-based synthetic circuits or biosensors (32,33). Timed supply of Acrs or genetic fusion of Acrs to Cas9 was further shown to reduce off-target editing in mammalian cells (13,34). Last but not least, we and others demonstrated a cell-type specific CasON switch based on microRNA dependent *acr* transgenes (11,12). This switch can restrict CRISPR-Cas activity to selected target cell types, e.g. hepatocytes or myocytes, and could aid focusing therapeutic genome editing to relevant tissues.

In all of the aforementioned applications, natural, constitutively active anti-CRISPR proteins were used. These can function exclusively as signaling mediators: To achieve a switch in CRISPR-Cas activity, the amount of Acr present in cells has to be controlled by means of supply or inducible expression. Toward unleashing the full potential of the Acr regulatory layer for CRISPR-Cas control, we previously developed the CASANOVA (**C**RISPR-Cas **a**ctivity **s**witching via **a n**ovel, **o**ptogenetic **v**ariant of **A**cr) concept, which facilitates switching off the activity of Acrs with blue light, thus releasing Cas9 activity (14,39). CASANOVA employs the light-oxygen-voltage 2 (LOV2) domain from *Avena sativa* (*As*) phototropin-1 for regulation of Acrs through photoreceptor insertion into allosteric, surface-exposed sites. This results in light-dependent CRISPR-Cas inhibitors that release Cas activity upon photoexcitation. We have previously demonstrated the CASANOVA concept for two different type II Acrs, namely AcrIIA4 and AcrIIC3, which are selective for *Spy*Cas9 and *Nme*Cas9, respectively. Subsequent studies reported slightly modified versions of our CASANOVA design employing a circularly permuted LOV2 receptor domain (40,41). Moreover, an interesting, recent study demonstrated that the fusion of CASANOVA directly into the PAM-interacting domain of Cas9 (AcrIIA4 acts as PAM mimic) results in single-chain, optogenetic Cas9-AcrIIA4-LOV2 hybrids (41). Beyond optogenetic Acr control, two previous studies reported chemically-inducible Acrs. In the first study, Song et al. created Acr-fusions to a 4-hydroxytamoxifen dependent intein. They employed several different Acrs, including AcrIIA4, -5, -25 and -32, the latter two of which were identified by the authors themselves (33). The system worked in principle for AcrIIA5, -26 and -32 and showed a noticeable rescue in Cas9 activity upon chemical induction. The dynamic range of control, however, was rather low, and for AcrIIA4, the applied control modality did not work at all (33). In the second study by Jain et al., the authors fused AcrIIA4 to a trimethoprim-dependent DHFR destabilization domain (DD), again resulting in trigger-dependent Acr activity and hence *Spy*Cas9 inhibition (42). Similar to the Song et al. approach, however, the DD-based strategy suffered from high leakiness and, as a result, a low dynamic range of control.

Therefore, with regard to Acr-based control of CRISPR-Cas systems with exogenous triggers, three central challenges remain. First, thus far, development of switchable Acr variants mainly focused on engineering Cas9 ortholog-specific Acrs that are not broadly applicable to likewise control many of the used Cas9 orthologs from all of the common type II subtypes (IIA, -B and -C). Second, the engineering of chemically-switchable Acr variants has thus far relied on ligand receptors and systems that lack overall potency, suffer from limited dynamic range, and are restricted to single drugs, which is suboptimal in terms of applicability and versatility for research use and potential clinical translation. Third, opto- and chemogenetic Acr control has thus far relied on a few Cas9-targeting inhibitors not compatible with other, widely used class II CRISPR-Cas types, such as the prominent type V/Cas12a clade.

Here, we present a switchable anti-CRISPR platform enabling multimodal control over a wide range of CRISPR-Cas orthologs across the type II and type V clades of class II CRISPR effectors. Building on our CASANOVA optogenetic strategy, we first engineered AcrIIA5 for light-dependent, broad-spectrum control of CRISPR-Cas9 activity via LOV2 domain insertion. Next, we expanded the CASANOVA approach to chemogenetic Cas9 control. Therefore, we developed a unique, versatile toolbox of circularly permuted human receptor domains that function as affinity clamps suitable for direct, allosteric Acr regulation. We show plug-and-play use of this novel chemoreceptor system for ligand-dependent activation of multiple type II Acrs, i.e. AcrIIA5, AcrIIA4, and AcrIIC3, and demonstrate modulation of Cas9 activity with six different, clinically relevant drugs. Finally, we extended both, optogenetic and chemical control, to type V CRISPR effectors by engineering light- and chemically switchable variants of AcrVA1. We believe that our toolbox holds great potential for regulating CRISPR-Cas9 and -Cas12 based systems in various contexts and opens new possibilities for engineering ligand-dependent, allosteric protein switches beyond the (anti-)CRISPR space.

## MATERIAL AND METHODS

### General cloning procedure

We used Golden Gate (43) and restriction enzyme cloning for construct assembly. Entry vectors encoding different Acrs were linearized by PCR for subsequent introduction of receptor domains (see below). Constructs used in this study are listed in Supplementary Table T1, and annotated sequence files are provided as Supplementary Data (GenBank files). Typically, oligonucleotides were ordered from Sigma-Aldrich. Genes for expression of different wild-type AcrIIA5 orthologs and AcrVA1 were obtained as synthetic double-stranded DNA fragments from Integrated DNA Technologies (IDT). For assembly of AcrIIA5-LOV2 and AcrVA1-LOV2 hybrids, the LOV2 domain and the respective Acr expression vector were PCR amplified with primers carrying 5’ overhangs introducing the Golden Gate sites, followed by Golden Gate cloning into PCR-linearized Acr-encoding vectors. To append linkers at Acr:LOV2 junction sites, LOV2 was PCR-amplified with primers carrying the linker-encoding sequences as 5’ overhangs, followed by Golden Gate assembly. DNA sequences encoding circularly permuted ligand receptors were obtained either as synthetic DNA fragement from IDT (for cpGR2 and UniRapR (44)) or N- and C-terminal parts were amplified in separate PCR reactions using constructs by Rihtar et al (45) obtained from Addgene (vectors were kind gifts from Roman Jerala; Addgene #188257, #188261, #188263, #188267) as templates (ERβ and TRβ). Amino acid sequences of cpReceptor domains are listed in Supplementary Table T2. These were then inserted at corresponding sites into AcrIIA5 and AcrVA1 encoding vectors using Golden Gate cloning as above. Expression vectors for CASANOVA-A4 and -C3 were reported by us in previous publications (14,39) and served as backbones for insertion of cpReceptor variants through Golden Gate assembly. The psiAAV dual luciferase reporter co-encoding firefly and *Renilla* luciferase was previously reported by us (14). We removed the *Renilla* gene from this vector by restriction cloning, resulting in a construct expressing firefly luciferase. A vector expressing *Renilla* luciferase only was obtained from Invitrogen (pRL-TK). The *Sau*Cas9 expression plasmid carrying a U6-driven crRNA scaffold and Golden Gate entry sites, i.e. plasmid pX601-AAV-CMV::NLS-SaCas9-NLS-3xHA-bGHpA;U6::BsaI-sgRNA, was a gift from Feng Zhang (Addgene plasmid # 61591 ; http://n2t.net/addgene:61591; RRID:Addgene_61591)(46). A vector co-expressing *Sau*Cas9 and a crRNA targeting the firefly luciferase gene (*Sau*Cas9_U6-sgRNA_FF1) was obtained by oligo cloning of a corresponding spacer sequence into this vector. SgRNAs and crRNAs were entered into corresponding U6-driven expression vectors by Golden Gate oligo cloning. Genomic target sites are listed in Supplementary Table T3. Pol-III promoter all-in-one CASANOVA-A5 constructs (Supplementary Figure S5), including those based on engineered, hybrid U6-H1 promoters (Supplementary Table T4), were obtained as synthetic DNA fragments and cloned into pcDNA3.1 carrying a sgRNA, 6×T-terminator, RBS and *Sau*Cas9-P2A-CASANOVA-A5 fragment, all assembled using restriction enzyme cloning. The vector encoding *Mb*Cas12a, named pCAG-hMb3Cas12a-NLS(nucleoplasmin)-3xHA (RTW2500), was a gift from Joseph Bondy-Denomy & Keith Joung & Benjamin Kleinstiver (Addgene plasmid # 115142 ; http://n2t.net/addgene:115142; RRID:Addgene_115142).

### Cell culture and transient transfection

Cell lines were maintained in a humidified incubator at 37°C with 5% CO₂ and were passaged every 3-4 days upon reaching 70–90% confluency. HEK293T (human embryonic kidney), HeLa (human cervical carcinoma), and HCT116 (colorectal cancer, kindly provided by Stefan Wölfl, University of Heidelberg) cells were cultured in phenol-red-free DMEM (Thermo Fisher/Gibco) supplemented with 10% (v/v) fetal calf serum (FCS, Biochrom AG), 2 mM L-glutamine, 100 U/ml penicillin and 100 µg/ml streptomycin (both from Thermo Fisher/Gibco). For transient transfection, cells were seeded in transparent, plastic 96-well plates (CytoOne) or black, glass-bottomed 96-well plates (Corning). 24 hours prior to transfection. HEK293T and HCT116 cells were seeded at a density of 12,500 cells in 100 µL medium per well, while HeLa cells were seeded at a density of 7,000 cells in 100 µL medium per well. HEK293T cells were transfected using Lipofectamine™ 2000 (Invitrogen, Thermo Fisher) and 0.5 µL Lipofectamine in a 50 µL transfection volume containing a total of 200 ng DNA per well. HCT116 and HeLa cells were transfected using Lipofectamine™ 3000 (Invitrogen, Thermo Fisher) with 0.3 µL Lipofectamine and 0.1 µL P3000 in a 15 µL transfection volume per well, also containing a total of 200 ng DNA and following the manufacturer’s protocol.

### Blue light setup

For optogenetic experiments, a previously reported LED device was used (47). Samples (well-plates) were illuminated within a cell culture incubator, using a custom-built blue light setup consisting of high-power blue LEDs powered by a Manson Switching Mode Power Supply (HCS-3102). The power supply was controlled by a Raspberry Pi running a custom Python script. Illumination was provided from below, through the transparent bottom of the 96-well plates, by positioning the plates on a custom-made sample holder with the LEDs mounted underneath. Samples were irradiated using a pulsatile illumination regime of 5 seconds on, 10 seconds off. The light intensity was approximately 5 W/m², as measured with a LI-COR LI-250A light meter.

### Dual luciferase reporter assay

HEK293T cells were seeded into black, clear-bottom 96-well plates (Corning) at a density of 12,500 cells per well in 100 µL of phenol-red-free DMEM (10% (v/v) FCS, 2 mM L-glutamine, and 100 U/ml penicillin plus 100 µg/ml streptomycin). 24 hours later, cells were co-transfected with (i) 40 ng of pSi-AAV, (ii) 20 ng of either *Sau*Cas9_U6-sgRNA_FF1 or control vector (CMV-*Sau*Cas9 with U6-sgRNA-scaffold and no spacer sequence), and (iii) 140 ng of the corresponding AcrIIA5-LOV2 hybrid, or CMV-AcrIIA5 or a DNA stuffer (as controls). For luciferase experiments with *Mb*Cas12a, we co-transfected cells with (i) 20 ng of a vector expressing firefly luciferase, (ii) 5 ng of a vector expressing *Renilla* luciferase, (iii) 20 ng of a vector encoding *Mb*Cas12a and a crRNA targeting the firefly luciferase gene, and (iv) 140 ng of the corresponding AcrVA1-GR2 hybrid, or wild-type AcrVA1 or DNA stuffer (as controls). Where applicable, cells were subjected to blue light illumination as described above. After 48 hours, firefly (FFLuc) and *Renilla* (Ren) luciferase activities were quantified using the Dual-Glo® Luciferase Assay System (Promega) following the manufacturer’s instructions. Cells were lysed in the supplied lysis buffer, and luminescence was measured using an Infinite M Plex microplate reader (TeCan). Measurements were taken with a 2-second delay after automatic substrate injection, followed by a 10-second integration time. Firefly luminescence was normalized to *Renilla* luminescence to correct for variations in cell number across wells.

### Cell viability Assay

Cell viability was assessed using the MTT Cell Proliferation Assay Kit (Cayman Chemical Company) following the manufacturer’s instructions. HEK293T cells were seeded into a 96-well plate at a density of 12,500 cells per well in 100 µL of phenol-red-free DMEM (10% (v/v) FCS, 2 mM L-glutamine, and 100 U/ml penicillin plus 100 µg/ml streptomycin). After an incubation time of 24 hours, respective drugs dissolved in DMSO were added to each well to reach the indicated final concentrations (Supplementary Figure S7 A). Following a 72 h incubation period, toxic Triton X-100 was added at different concentrations ranging from 1% to 0.001% to untreated wells as a positive control for cell death. MTT treatment was initiated by adding 10 µL of MTT reagent to each well, followed by a 3–4 hour of incubation. Subsequently, 100 µL of freshly prepared crystal-dissolving dye was added to each well, and the plate was incubated overnight for 16-18 h to ensure complete solubilization. Absorbance was measured at 570 nm using an Infinite M Plex microplate reader (TeCan). Untreated cells served as a control for maximum cell viability.

### T7E1 assay and NGS sequencing

Editing efficiency (%) was assessed via the T7 Endonuclease1 (T7E1) assay or targeted amplicon sequencing. A list of all genomic target loci and primers used for PCR amplification and NGS sequencing is available in Supplementary Table T5.

HEK293T, HeLa, and HCT116 cells were seeded in 96-well plates 24 hours prior to transfection. Tansfections were performed using Lipofectamine™ 2000 (Invitrogen, Thermo Fisher) for HEK293T cells and Lipofectamine™ 3000 (Invitrogen, Thermo Fisher) for HeLa and HCT116 cells, with a total DNA amount of 200 ng per well.

For experiments in Figure 1 E and F, (i) 100 ng of *Sau*Cas9-sgRNA expression vector and (ii) 50 ng of CASANOVA-A5 or wild-type AcrIIA5 (as control) were co-transfected. For experiments in Supplementary Figure S5, 200 ng of an all-in-one Pol-III driven sgRNA-*Sau*Cas9-CASNOVA-A5 vector was transfected. A corresponding vector without AcrIIA5 was used as control. For experiments with cpReceptor-based AcrIIA5 variants (CASANDRA), 100 ng of *Sau*Cas9-sgRNA was co-transfected with 10 ng of vectors encoding AcrIIA5-cpReceptor hybrids or wild-type AcrIIA5 (as control), unless otherwise specified in the figures. For the use of CASANDRA-A5 to control other orthologs than *Sau*Cas9, we co-transfected either (i) 100 ng of *Nme*Cas9-sgRNA vector with 50 ng of CASANDRA-A5 or wild-type AcrIIA5 vector (as control) or (ii) 60 ng of *Spy*Cas9 vector with 15 ng of sgRNA vector and 50 ng of CASANDRA-A5 or wild-type AcrIIA5 vector (as control). For experiments with AcrIIC3-GR2/A4-GR2 hybrids (CASANDRA-C3, CASANDRA-A4), we co-transfected either (i) 150 ng of *Nme*Cas9-sgRNA and 10 ng of CASANDRA-C3 or wild-type AcrIIC3 (as control) or (ii) 33 ng of each, the *Spy*Cas9 and sgRNA expression vectors and 90 ng of CASANDRA-A4 or wild-type AcrIIA4 (as control). Inhibition of *Sau*Cas9 by AcrIIA5-GR2 in HeLa and HCT116 cells was performed by co-transfecting 150 ng of *Sau*Cas9 vector and 10 ng of AcrIIA5-GR2 or wild-type AcrIIA5 vector (as control). For experiment with *Mb*Cas12a, we co-transfected 30 ng of *Mb*Cas12a vector, 30 ng of crRNA expression vector and 140 ng of AcrVA1-GR2 hybrid or wild-type AcrVA1 (control) encoding vector. Total DNA amounts per well were topped up to 200 ng with a irrelevant stuffer plasmid (pBluescript) where needed. Non-targeting sgRNA/crRNA control vectors lacked the corresponding spacer sequences.

**Figure 1.**
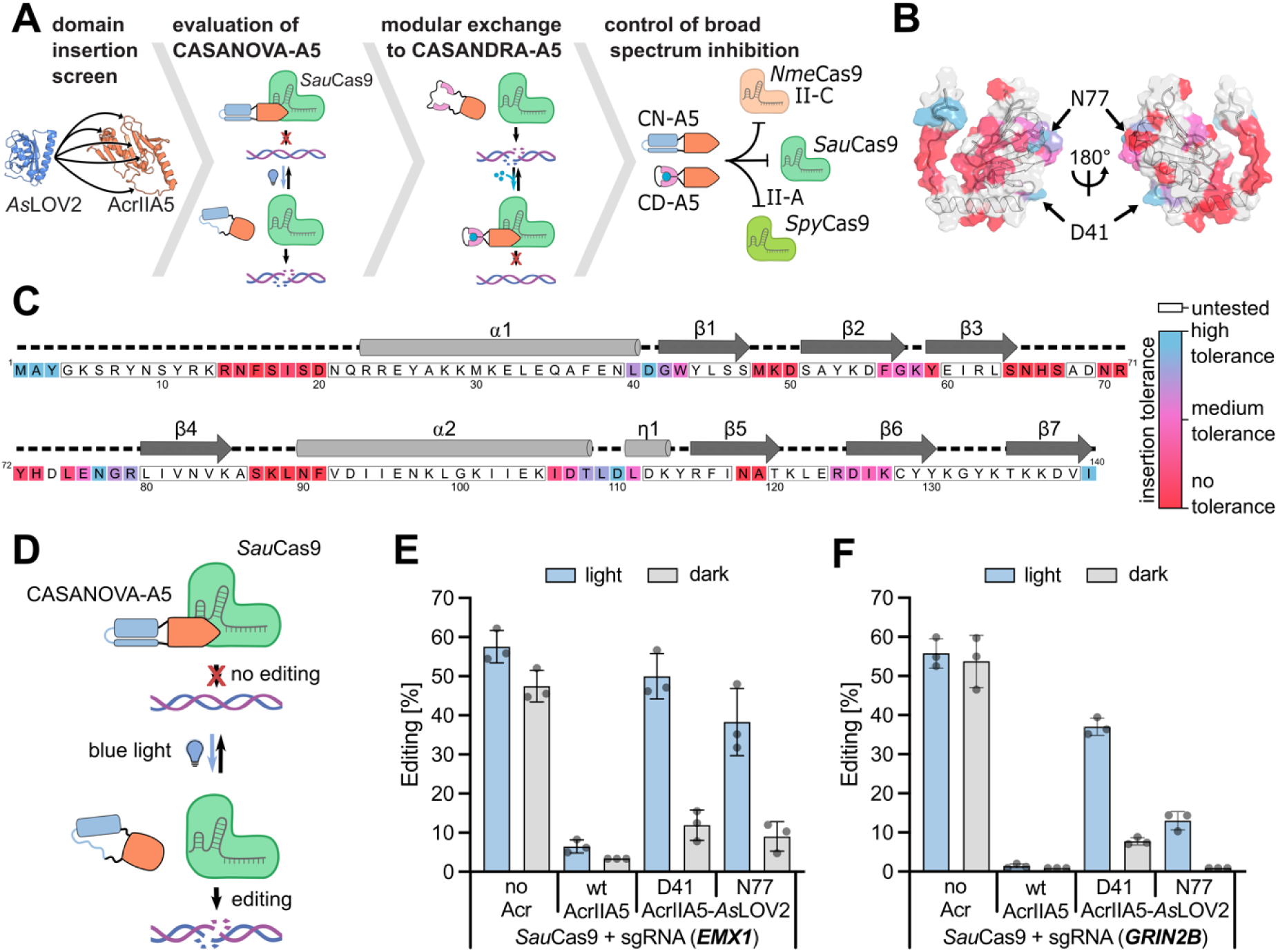
Engineering optogenetic control of AcrIIA5 by LOV2 domain insertion. (A) Schematic indicating the process for engineering switchable variants of AcrIIA5. (B) Structure of AcrIIA5 (PDB ID: 2zva). The color coding indicates LOV2 domain insertion tolerance as measured by luciferase assay (refer to Supplementary Figure S3). Positions in red do not tolerate LOV2 domain fusion preceding the indicated residue, blue indicates positions tolerating domain fusion. White indicates positions that were not experimentally assessed. (C) Primary amino acid sequence and corresponding secondary structure elements (according to (37)). Color code as in B. (D) Schematic of CASANOVA-A5 function. (E, F) HEK293T cells were co-transfected with plasmids encoding the indicated components and illuminated for 72 h or kept in the dark, followed by NGS analysis of InDel frequencies at the *EMX1* (E) and *GRIN2B* (F) locus. Bars indicate means, grey dots individual data points, and error bars the SD from n = 3 independent experiments.

Where applicable, chemicals dissolved in DMSO were added 2 hours post-transfection, while for blue light induction experiments, light exposure of samples started 6 hours after transfection, with dark control samples kept in the dark throughout the entire experiment. No ligand control samples were treated with identical amounts of DMSO. Indicated concentrations of chemical ligands were calculated for a working volume per well of 100 µL (pre-transfection). Following a 72 hour incubation post transfection, media was discarded and cells were lysed in 141.4 µL of DirectPCR lysis reagent (Peqlab) supplemented with proteinase K (Sigma) and incubated at 55°C for 6 hours at 120 rpm. Proteinase K was inactivated by heat treatment at 85°C for 45 minutes. Genomic DNA was then PCR-amplified using the Q5® High-Fidelity 2X Master Mix (NEB) and primers flanking the target site carrying Illumina-compatible 5’ adaptors and barcodes (Supplementary Table T5) for multiplexing during NGS.

To prepare samples for NGS analysis, PCR amplicon quality was assessed by running samples on a 1% Tris-acetate-EDTA (TAE) agarose gel in 0.5x TAE buffer and DNA bands were quantified using the gel analysis tool in ImageJ (version 1.54k; https://imagej.net). For multiplexing, barcoded PCR amplicons were combined by mixing at equimolar quantities, followed by purification using the QIAquick PCR Purification Kit (QIAGEN). DNA concentration was measured with a NanoPhotometer® NP80 (IMPLEN) and diluted to 20 ng/µL in a total volume of 25 µL. Illumina sequencing was performed using the Amplicon-EZ service (GeneWiz) and editing percentages (Editing (%)) were calculated by demultiplexing samples based on barcodes, followed by InDel quantification with CRISPResso2 (48). To assess the Illumina sequencing errors arising due to multiplexing (i.e. index swapping between edited and no-edited samples), a separate sequencing run with unique, double-indexing of control samples was performed to identify the actual leakiness of wild-type Acrs with respect to Cas effector inhibition (Supplementary Figure S1). Difference in the resulting values and the corresponding background values in the data were considered index swapping errors and substracted from respective samples.

To perform T7 Endonuclease I (T7E1) assay, heteroduplex DNA was generated by diluting 5 µL of PCR amplicon in 20 µL of 1x NEB buffer 2, followed by heating to 95°C for 5 minutes, and gradually cooling down samples to 25°C in a nexus gradient GSX1 Mastercycler (Eppendorf). Afterward, 0.5 µL of T7 endonuclease I (NEB) was added, samples were mixed, and incubated at 37°C for 15 minutes. Products were analyzed on a 2% Tris-borate-EDTA (TBE) agarose gel in 1× TBE buffer, followed by gel image acquisition using a GelDoc system equipped with a 2.8-megapixel, 14-bit scientific-grade CCD camera (Intas). Editing efficiency was quantified from gel images by background subtraction and peak area analysis using the ImageJ gel analysis tool (version 1.54k; https://imagej.net). Editing percentage (Editing (%)) was calculated as Editing (%) = 100 × (1 – (1 – Fraction cleaved)^1/2), where Fraction cleaved = (∑Cleavage product bands)/(∑Cleavage product bands + PCR input band).

### Protein structure prediction and visualization

Protein structures were predicted with AlphaFold 3 (49) (as of Nov. 2024; https://alphafoldserver.com/about) with standard settings, followed by color-coding and rendering in PyMOL (version 3.0.0; https://www.pymol.org).

### Statistics

Independent experiments refer to samples prepared, treated and measured independently on different days, starting with separate cell cultures. For T7E1 and NGS data sets, bars indicate means, individual data points (grey dots) indicate data from different days, error bars depict the standard deviation (SD). For luciferase experiments and the MTT assay, grey dots indicate individual technical replicates (3 technical replicates correspond to one experiment), bars are means of all technical replicates and error bars correspond to the SD. No samples were excluded from the analysis.

### English language editing

ChatGPT version 4.o (OpenAI) was used for English language editing.

## RESULTS

### AcrIIA5 tolerates *As*LOV2 insertion at various surface sites, which enables optogenetic control of *S. aureus* Cas9

Towards engineering light-switchable AcrIIA5 variants (Figure 1A), we first compared the inhibition potency of different AcrIIA5 orthologs in human cells. Therefore, we co-expressed AcrIIA5 derived from *Streptococcus thermophilus* (*S. th.*), *Streptococcus pyogenes* strain D4276, *Enterococcus faecalis*, *Dolosigranulum pigrum*, *Lactobacillus sakei*, *Streptococcus pneumoniae* strain P4761 (*S. pneu.* P4761), and *Streptococcus mitis* bacteriophages with either *Sau*Cas9 or *Spy*Cas9 in HEK293T cells alongside sgRNAs targeting the *EMX1* or *CCR5* locus, respectively. Using T7 endonuclease I (T7E1) assay, we then qualitatively assessed editing efficacy at the targeted genomic sites. We observed highly potent inhibition of both, *Sau*Cas9 and *Spy*Cas9, for most AcrIIA5 orthologs except for the variant derived from *Streptococcus pneumoniae* strain P4761 bacteriophage (Supplementary Figure S2). Hence, we decided to proceed with the *S. th.* ortholog, since it is well-characterized and its structure has been resolved (26,28,37,50).

Next, we aimed at identifying surface sites in AcrIIA5 amenable to domain fusion and hence potentially suited to create allosteric, light-switchable AcrIIA5 derivatives through *As*LOV2 insertion. On the basis of the reported AcrIIA5 structure (37), we identified several potential candidate sites located mainly in loop regions exposed on the protein’s surface (Figure 1B). Using these sites as seed points, we then inserted the LOV2 domain at several positions located around these sites in the primary sequence (Figure 1C). Subsequently, we assessed the ability of the resulting AcrIIA5-LOV2 hybrids to inhibit the *Sau*Cas9-mediated cleavage of a luciferase reporter in HEK293T cells following expression for 48 h in the dark. Note, that we performed this initial screen in the absence of light, since the LOV2 takes on a compact conformation under this condition, which should, ideally, not perturb the conformation of the fused AcrIIA5 part and hence facilitate the identification of hybrid variants that generally retain activity. Remarkably, various AcrIIA5-LOV2 hybrids showed potent inhibition of *Sau*Cas9, evidenced by a strong rescue in luciferase activity (Figure 1B,C and Supplementary Figure S3). Active hybrids contained the LOV2 insertions in various parts of AcrIIA5, both on the primary sequence and structural level, i.e. at residues 40-43, 56-69, 75-79, 106-111, 124-127, as well as the protein’s N- and C-terminal parts (Figure 1B,C and Supplementary Figure S3). This was rather unexpected, since our previous domain insertion studies in context of AcrIIA4 and AcrIIC3 typically yielded very few positions that tolerated domain fusion (14,39). In contrast, AcrIIA5 appears to be rather promiscuous with respect to domain insertion tolerance.

Next, we selected two highly active insertion variants carrying the LOV2 domain in front of residues D41 or N77 in AcrIIA5 for subsequent validation of light-dependent *Sau*Cas9 inhibition and hence optogenetic genome editing (Figure 1D). To this end, we co-transfected HEK293T cells with constructs (i) encoding *Sau*Cas9 and a sgRNA targeting the *EMX1* or *GRIN2B* loci as well as (ii) the respective AcrIIA5-LOV2 fusion. Following incubation of cells under pulsatile blue light or in the dark for 72 h, InDel frequencies were assess by NGS. Excitingly, for both AcrIIA5-LOV2 hybrids, we observed potent, light-induced genome editing on both loci and low InDel frequencies in the absence of light (Figure 1E and F). Control samples expressing no Acr or wild-type AcrIIA5 exhibited high or low editing irrespective of illumination, respectively (Figure 1E and F). In continuation with our previously reported, optogenetic Acr derivatives (14,39), we named these AcrIIA5-LOV2 hybrids CASANOVA-A5 (for **C**RISPR-Cas **a**ctivity **s**witching via **a n**ovel **o**ptogenetic **v**ariant of **A**crIIA5). Of note, the CASANOVA-A5 based on the D41 insertion site showed, overall, higher background editing in the system’s dark state (off-state) as compared to the N77 variant, but facilitated release of Cas9 activity to levels similar to the positive control in the light-induced state (on-state; Figure 1E and F). In turn, the N77 variant was less leaky in the dark, but also compromised Cas9 activity in the illuminated state to a noticeable extend (Figure 1E and F; see Discussion).

Thus far, we have observed effects resulting from a naïve insertion of LOV2 into AcrIIA5. From our previous work (14) as well as literature (51,52), however, it is well known that the residues linking the receptor and effector parts of hybrid proteins can have a strong influence on the performance of allosteric protein switches. Thus, we performed a small screen and appended symmetric linkers of different length (2-12 residues) and composition to the LOV2:Acr junctions in CASANOVA-A5. These ranged from GS-rich flexible linkers, through semi-rigid linkers (e.g. PAS and TPT linkers) to rigid proline-based linkers (Supplementary Figure S4). As expected, the linker length and composition had a strong impact on CASANOVA-A5 performance (Supplementary Figure S4). Most linkers increased inhibition potency of CASANOVA-A5 in the dark. This favorable property, however, came at the cost of overall compromised activity of Cas9 in the on-state due to some inhibitory activity being still present upon photoexcitation. This indicates an intrinsic compromise between the system’s leakiness in the dark and its ability to effectively release and hence render active Cas9 upon light-mediated activation (see Discussion).

A particularly attractive trait of CASANOVA-A5 is its small size of only 283 amino acids. At the same time, *Sau*Cas9 is also among the smaller Cas9 orthologs with about 1054 residues in length. We reasoned that it may be possible to co-encode CASANOVA-A5 as well as Cas9 and a sgRNA as a single transgene that would remain within the size-limit for Adeno-associated viral (AAV) vectors (∼4.8 kbp), a highly promising gene delivery vehicle (53,54). To explore if we could “squeeze” our entire system to AAV-compatible size, we adapted a previously-reported co-expression strategy hijacking the polymerase-II (pol-II) co-activity of pol-III promoters (55). Pol-III promoters commonly used in CRISPR for sgRNA expression, such as U6 and H1, show certain levels of pol-II activity, and the pol-II/pol-III activities can be tuned by promoter engineering (56,57). Building on this concept, we speculated that pol-III promoters could not only be used for co-expression of Cas9 and a sgRNA, as previously already shown, but via a 2A-peptide strategy could enable additional co-expression of further proteins, such as our CASANOVA-A5.

To explore this concept, we created constructs encoding (i) an upstream H1 or U6 promoter or synthetic hybrids thereof engineered by us, followed by (ii) a sgRNA coding sequence (spacer, scaffold, 6×T pol-III terminator), which was followed by (iii) a strong RBS as well as a sequence encoding *Sau*Cas9-P2A-CASANOVA-A5 and a pol-II terminator (Supplementary Figure S5 A). This strategy yielded sizes of the entire expression cassette between 4.45 kbp (for H1-based variants) to ∼4.6 kbp (for U6-based variants) co-encoding all components required for optogenetic control of *Sau*Cas9, which is well within the AAV cargo size limit. We transfected these variants into HEK293T cells (targeting either *EMX1* or *GRIN2B*), exposed cells to blue light for 72 h or kept them in the dark, followed by assessment of editing using T7E1 assay. As hoped, we observed potent, light-dependent CRISPR editing for several constructs on both loci (Supplementary Figure S5 B,C). H1-based variants encoding the CASANOVA-A5 variant based on the N77 insertion site were particularly promising and showed robust, light-dependent editing on both targeted loci. We conclude that co-expression of all three components, Cas9, sgRNA and CASANOVA-A5 is in principle possible from a single transgene fitting within the AAV cargo limit (see Discussion).

### Engineering circularly permuted human receptor domains for ligand-dependent AcrIIA5 activation

The CASANOVA concept has thus far been limited to the LOV2 domain and hence to light-dependent control of Acrs. We envisioned that a similar concept could also be implemented on the basis of ligand-responsive receptor domains, transferring the CASANOVA concept to chemical Acr control. Mammalian hormone receptor domains have been widely adopted in synthetic biology for controlling the activity of proteins of interest. Applying hormone receptors for allosteric control via receptor domain insertion into an effector protein is, however, challenging, since usually, the N- and C-termini in these receptors are located relatively far apart. Thus, receptor insertion into the allosteric surface site of an effector protein is likely to perturb the protein’s structure and hence effector activity, at least to some extent. For example, fusing an estradiol receptor domain into an allosteric surface site of CRISPR-Cas9 yielded Cas9-receptor hybrids that, while dependent on 4-OHT, showed a strongly diminished overall activity as compared to wild-type Cas9 (18).

Recently, Rihtar et al. reported split variants of several human hormone and glucocorticoid receptor domains that respond to clinically-relevant drugs and can be used to effectively implement split-protein control strategies in mammalian cells (45). We reasoned that these receptors, with their identified split-sites, would be ideal starting points for engineering circularly permuted human receptor domains that function as drug-dependent “affinity clamps” well-suitable for allosteric control of Acrs (Figure 2A) and, potentially, other effector proteins (see Discussion). To implement this novel concept, we started from three native receptor domains, namely the estradiol receptor β (ERβ) activated by β-estradiol (β-EST) and 4-hydrotamoxifen (4-OHT), the thyroid receptor β (TRβ) responding to triiodothyronine (T3) as well as the glucocorticoid receptor 2 (GR2) controlled by cortisol (COR), dexamethasone (DEX) and mometasone furoate (MOF) (Figure 2B and Supplementary Figure S6). We chose these three receptors, since they respond to various, clinically-approved drugs that are well-tolerated in mammalian cells over a wide concentration range (Supplementary Figure S7 A). Also, these drugs do not affect CRISPR genome editing efficacy when applied at reasonable concentrations (Supplementary Figure S7 B-G).

**Figure 2.**
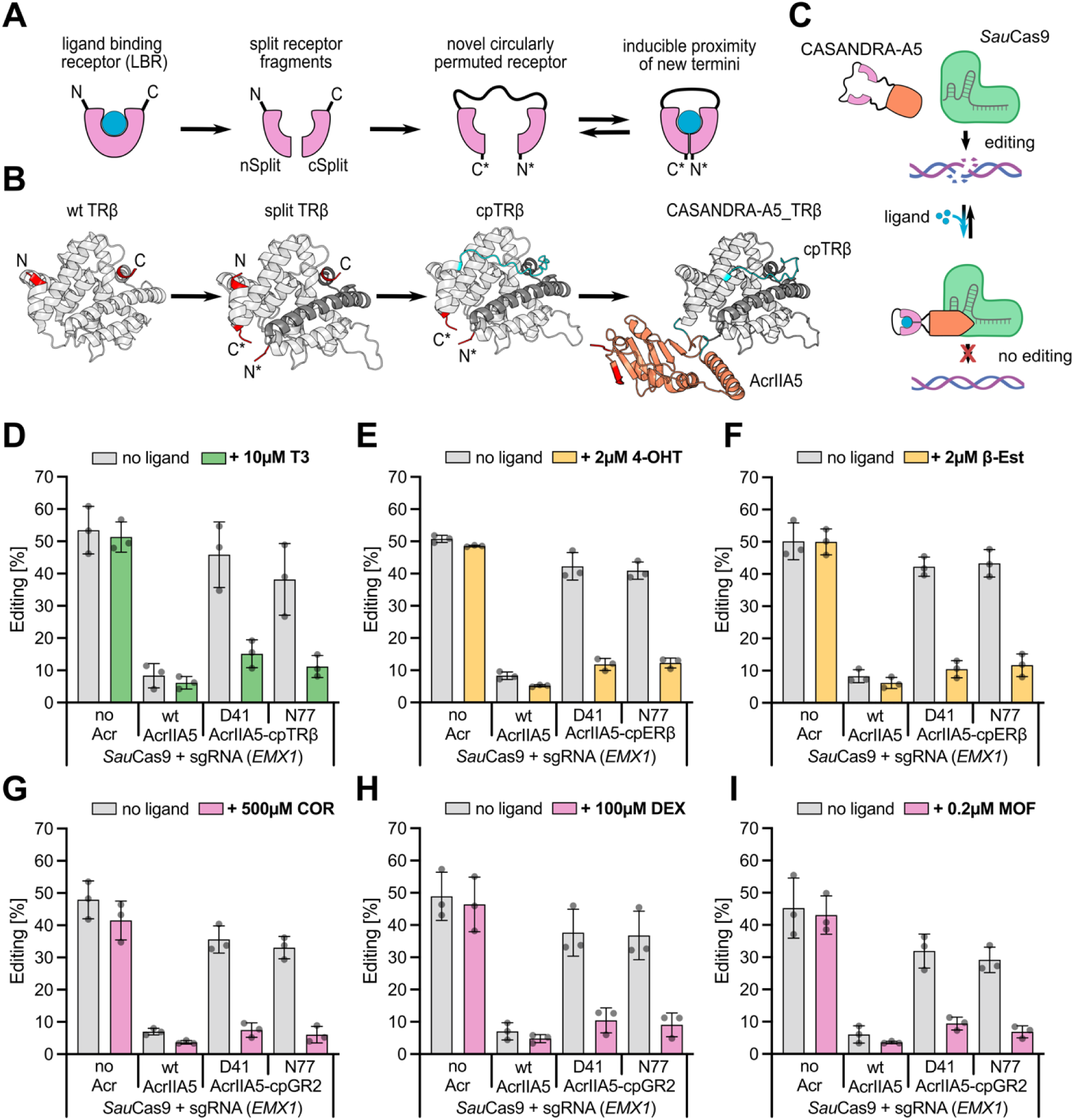
Design and application of circularly permuted receptor domains for ligand-dependent activation of AcrIIA5. (A) Schematic showing the general design strategy for engineering cpReceptor variants. Receptors were opened at previously identified split-sites (45), thereby creating new N- and C-termini that are in direct proximity (C*, N*). Former N- and C-termini were connected by fusion via a long GS-linker (black). Resulting cpReceptors should take on a compact conformation, with proximate N- and C-termini, upon chemical induction (blue). (B) In analogy to the schematic in A, cpReceptor engineering stages are shown for the thyroid hormone receptor beta domain (TRβ; PDB ID: 1xzx; first three structures from the left). The structure on the right shows an AlphaFold3 prediction of the fusion between the resulting cpTRβ domain inserted at D41 into AcrIIA5. red: termini; blue: linker residues. (C) Schematic of CASANDRA-A5 function. (D-I) HEK293T cells were co-transfected with the indicated components and treated with the indicated drugs starting 2 h later. 72 h post transfection InDels at the targeted *EMX1* locus were analyzed by NGS. Bars indicate means, grey dots individual data points, and error bars the SD from n = 3 independent experiments. The specific cpReceptor:ligand pairs are as follows. (D) cpTRβ:triiodothyronine (T3). (E) cpERβ:4-hydroxytamoxifen (4-OHT). (F) cpERβ:β-estradiol (β- EST). (G) cpGR2:cortisol (COR). (H) cpGR2:dexamethasone (DEX). (I)cpGR2:mometasone furoate (MOF).

We then re-located the receptor’s termini by opening up the proteins at the previously identified split-sites, resulting in new N- and C-termini positioned in direct spatial proximity (Figure 2B and Supplementary Figure 6). The former N- and C-termini were, in turn, connected by long, flexible linkers, yielding single-chain, circularly permuted (cp) receptor domains, which due to the proximity of the new termini, should be ideally suited for protein control via receptor insertion into allosteric surface sites (Figure 2B and Supplementary Figure 6). We then fused the cp receptors into AcrIIA5 (at positions preceding D41 or N77), yielding AcrIIA5-cpReceptor fusions expected to mediate effective, broad-spectrum CRISPR-Cas9 inhibition in presence of their corresponding ligand. To identify a suitable working dose of the AcrIIA5-cpReceptor constructs for transient transfection experiments, we first performed a Cas9 inhibition experiment in absence of the ligands using the cpTRβ and cpERβ variants. Therefore, we titrated the amount of AcrIIA5-cpReceptor encoding plasmid between 100 ng and 10 ng (per well in 96-well format) during transfection into HEK293T cells, while keeping the amount of *Sau*Cas9/sgRNA vector constant (100 ng), followed by T7E1 assay. At low (10 ng) to medium (33 ng) plasmid doses, the AcrIIA5-cpReceptor hybrids did not affect or only mildly affected Cas9 activity (as intended), while at very high doses (100 ng), considerable, unintended CRISPR-Cas9 inhibition was observed (Supplementary Figure S8). We hence selected the 10 ng and 33 ng plasmid dose for subsequent genome editing experiments in HEK293T cells, this time adding the corresponding ligands at pre-selected, non-toxic concentrations (Supplementary Figure S7). We observed potent, ligand-dependent genome editing in HEK293T cells at the *Sau*Cas-targeted *EMX1* locus for all AcrIIA5-cpRecptor:ligand pairs by T7E1 assay (Supplementary Figure S9 and S10) and NGS (Figure 2 D-I). Finally, we compared the efficiency of our engineered cpReceptors to UniRapR (44), a previously reported, rapamycin-dependent affinity clamp well-known in synthetic biology. In analogy to our AcrIIA5-cpReceptor hybrids, we fused the UniRapR domain at the identified positions (D41 or N77) into AcrIIA5 and assessed rapamycin-dependent *Sau*Cas9 inhibition in HEK293T cells. Our cpReceptor based AcrIIA5 hybrids clearly outperformed the UniRapR-based hybrids, showing much lower genome editing background in presence of the ligand and potent rescue of editing close to the positive control levels in absence of the ligand (Supplementary Figure S10; compare panels A-F with panel G).

We refer to our Acr-cpReceptor hybrids as **CASANDRA: C**hemical **A**ctivation of **S**witchable **A**nti-CRISPRs via **N**ovel **D**rug-responsive **R**eceptor **A**rchitectures. In analogy to CASANOVA, the AcrIIA5-based CASANDRA variants are referred to as CASANDRA-A5.

### CASANDRA-A5 is a ligand-activated, broad spectrum CRISPR-Cas9 inhibitor and functions in different human cell lines

Due to the broad-spectrum activity of AcrIIA5, it is to be expected that CASANDRA-A5 can be applied for ligand dependent-control not only for *Sau*Cas9, but also other prominent Cas9 orthologs. To demonstrate this, we co-expressed the CASANDRA-A5 variants based on the cpERβ or cpGR2 domains with (i) *Nme*Cas9 and a sgRNA targeting the *F8* locus or (ii) *Spy*Cas9 and a sgRNA targeting the *CCR5* locus in HEK293T cells (Figure 3A and D). We then incubated cells in presence or absence of corresponding ligands (4-OHT or COR) for 72 h, followed by NGS. This resulted in potent, ligand-dependent inhibition of both CRISPR-Cas orthologs, while positive (no Acr) and negative (wild-type AcrIIA5) control samples showed high or low editing irrespective of ligand addition (Figure 3B,C and E,F).

**Figure 3.**
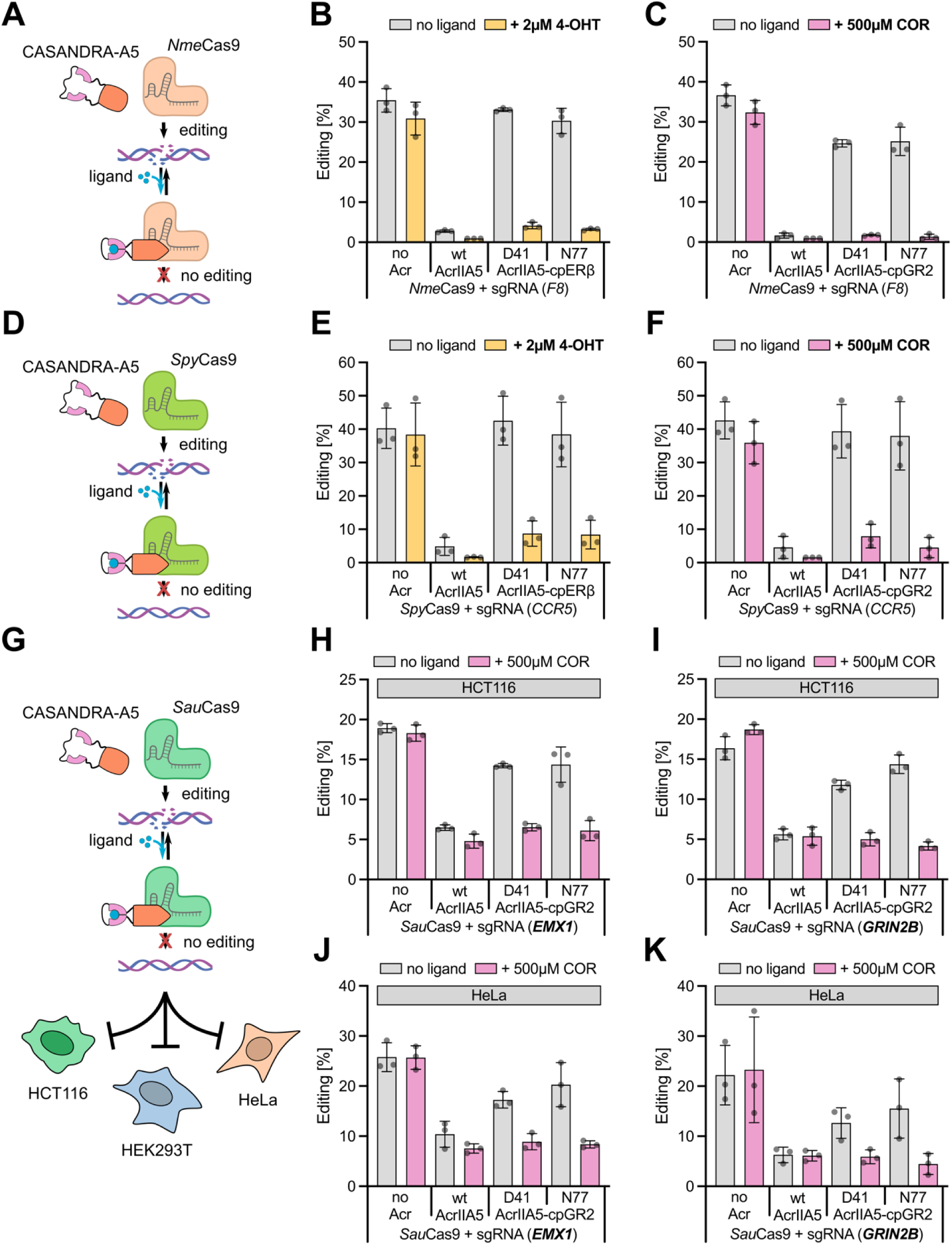
CASANDRA-A5 functions as a switchable, broad-spectrum inhibitor and can be applied in different human cell lines. (A, D) Schematic of CASANDRA-A5 mediated control of *Nme*Cas9 (A) and *Spy*Cas9 (D). (B, C, E, F) HEK293T cells were co-transfected with plasmids encoding (i) *Nme*Cas9 and a sgRNA targeting the *F8* locus (B, C) or *Spy*Cas9 and a sgRNA targeting the *CCR5* locus (E, F) and (ii) the indicated AcrIIA5-cpReceptor hybrid. Cells were treated with the indicated drugs starting 2 h post transfection and incubated for an additional 70 h, followed by NGS analysis of InDels at the targeted locus. Bars indicate means, grey dots individual data points, and error bars the SD from n = 3 independent experiments. The specific cpReceptor:ligand pairs are as follows. (B, E) cpERβ:4-OHT. (C, F) cpGR2:COR. (G) Schematic of CASANDRA-A5 mediated *Sau*Cas9 control in different cell lines. (H-K) HCT116 (H, I) or HeLa (K, J) cells co-expressing the indicated components were treated with COR as in C. 72 h post transfection, cells were lysed, followed by NGS analysis of InDels at the targeted *EMX1* (H, J) or *GRIN2B* (I, K) locus. Bars indicate means, grey dots individual data points, and error bars the SD from n = 3 independent experiments.

Subsequently, we tested if CASANDRA-A5 was suitable for CRISPR-Cas9 control in different cells lines (Figure 3G). To this end, we co-expressed the CASANDRA-A5 variant based on cpGR2 with *Sau*Cas9 and a sgRNA targeting either *EMX1* or *GRIN2B* in HCT116, a human colon cancer cell line, or HeLa, a human cervix carcinoma lines in presence or absence of the corresponding ligands for 72 h, followed by NGS. Again, we observed ligand-dependent inhibition of Cas9 activity (Figure 3H-K). Of note, the overall editing efficacy (positive control) was lower and background editing in presence of the wild-type AcrIIA5 control higher as compared to HEK293T cells (compare Figure 3B,C,E, F to H-K). This is likely due to overall lower transfection efficacy and thus uneven expression of Cas9:CASANDRA:sgRNA components in individual cells (see Discussion). In sum, these experiments evidence the great potential of CASANDRA for ligand-dependent regulation of CRISPR-Cas9 activity in human cells.

### Chemogenetic control with engineered cpReceptor domains can be plug-and-play transferred to other anti-CRISPR proteins

Encouraged by the robust functionality of our engineered cpReceptors, we aimed to extend the CASANDRA approach to other anti-CRISPR proteins. This effort was driven by two key motivations. First, a successful adaptation would create a modular toolbox of ligand-responsive Acrs, enabling precise control over or enable fine-tuning of diverse CRISPR-Cas orthologs in human cells. This is particularly valuable since different Acrs vary significantly in their target specificity, inhibition potency, and mechanisms of action against CRISPR-Cas systems. Second, the structural diversity of Acrs provides an excellent test case to evaluate the robustness of our cpReceptor design for allosteric protein control via receptor insertion.

Previously, we reported optogenetic derivatives of AcrIIA4 and AcrIIC3 created via LOV2 domain fusion. For AcrIIA4, the LOV2 domain was inserted between residues W63 and Y67 (with residues 64–66 being removed), which yielded CASANOVA-A4, an optogenetic inhibitor of *Spy*Cas9. Similarly, in AcrIIC3, the LOV2 was inserted after residue F59, yielding a light-dependent inhibitor of *Nme*Cas9.

We reasoned that the LOV2 domain in CASANOVA-A4 and -C3 could be replaced with cpReceptor domains in a plug-and-play fashion. To investigate this, we created CASANDRA-A4 and CASANDRA-C3 through insertion of the GR2 domain into the aforementioned insertion sites. We then co-expressed the CASANDRA variants with their cognate Cas9 ortholog, i.e. *Spy*Cas9 or *Nme*Cas9, as well as a sgRNA targeting a genomic site in HEK293T cells (Figure 4A and C). We observed potent, ligand-dependent inhibition of both CRISPR-Cas orthologs (Figure 4B and D), indicating that the cpGR2 receptor facilitates chemogenomic control of various type II Acrs.

**Figure 4.**
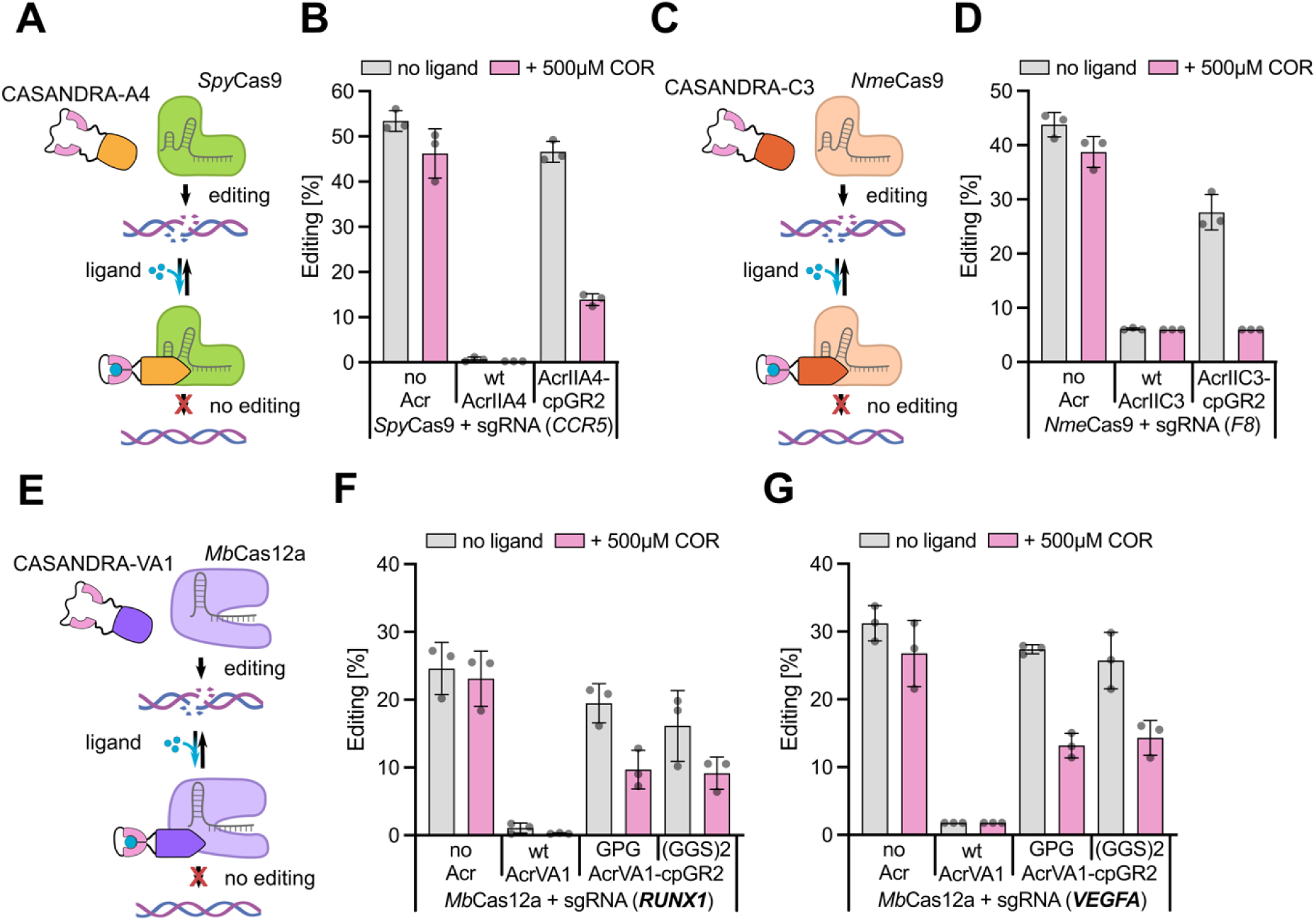
Chemogenetic control of other type II anti-CRISPR proteins and fine-tuning of Cas12a through ligand-responsive AcrVA1. (A, C) Schematic of CASANDRA-A4 mediated control of *Spy*Cas9 (A) and CASANDRA-C3 mediated control of *Nme*Cas9 (C). (B,D) HEK293T cells were co-transfected with the indicated components and treated with COR starting 2 h later. 72 h post transfection, cells were lysed, followed by NGS analysis of InDels at the targeted locus (indicated). Bars indicate means, grey dots individual data points, and error bars the SD from n = 3 independent experiments. (E) Schematic of CASANDRA-VA1 mediated control of *Mb*Cas12a. (F, G) HEK293T cells were co-transfected with the indicated components and treated with COR starting 2 h later. 72 h post transfection, cells were lysed, followed by NGS analysis of InDels at the targeted *RUNX1* (F) or *VEGFA* (G) locus. Bars indicate means, grey dots individual data points, and error bars the SD from n = 3 independent experiments.

### Transferring the CASANOVA and CASANDRA concept to AcrVA1 for fine-tuning of Cas12a

Finally, we sought out to explore, if our opto- and chemogenetic Acr control strategies could be translated to CRISPR-Cas orthologs other than Cas9. AcrVA1 is a broad-spectrum inhibitor of CRISPR-Cas12a. Since no allosteric surface sites were known in AcrVA1 yet, we assessed the previously reported AcrVA1 structure in complex with Cas12a (58) for potential candidate sites (Supplementary Figure S11 A). A surface exposed stretch of residues carrying a small helical element (residues S103 to I109) appeared promising, since it extruded away from the Cas12a/AcrVA1 complex and hence, fusion of an additional domain should not cause steric clashes with Cas12a. We sampled this region residue-wise through insertion of LOV2. On top, we systematically deleted AcrVA1 residues around the insertion site to facilitate tight embedding of the receptor domain into the AcrVA1 structure, akin to our previous, successful attempts for AcrIIA4 (14). We then assessed the activity of the resulting AcrVA1-LOV2 hybrids, first in the absence of light, using a luciferase reporter assay, in which *Mb*Cas12a targets and cleaves a firefly luciferase gene. Several AcrVA1-LOV2 hybrids showed a mild, but noticeable inhibition of Cas12a as evidenced by a rescue in firefly luciferase activity (Supplementary Figure S11B). Next, we followed up on one promising variant bearing the LOV2 domain between AcrVA1 residues 103 and 107, i.e. the construct contains a 3-residue deletion, which roughly corresponds to a single alpha helix turn. This variant was further modified by appending flexible, symmetric GS-linkers of different length at both LOV2-AcrVA1 junction sites. We again assessed the activity of the resulting, second-generation AcrVA1-LOV2 hybrids using the aforementioned luciferase assay, this time incubating cells either under pulsatile blue light or in the dark for 48 h prior measurement. Several hybrid inhibitors showed a promising, light-dependent activity, resulting in marked Cas12a inhibition in the dark and complete recovery of Cas12a activity upon illumination (Supplementary Figure 11 C). Of note, inhibition in the dark was still lower than that observed for wild-type AcrVA1, indicating that these variants may still hold potential for further improvement by engineering (see Discussion). The AcrVA1-LOV2 hybrid bearing a 4-residue flexible linker was the most potent, hybrid inhibitor and showed strong light-dependent Cas12a release (Supplementary Figure 11C and D). We named this variant CASANOVA-VA1.

Finally, we replaced the LOV2 domain in CASANOVA-VA1 with the cpGR2 receptor to assess the possibility of ligand-dependent tuning of Cas12a activity in human cells. We co-expressed the resulting CASANDRA-VA1 with *Mb*Cas12a as well as crRNAs targeting the *RUNX1* or *VEGFA* loci in HEK293T cells (Figure 4E). Cells were then incubated in presence or absence of COR for 72 h, followed by NGS. We reproducibly observed ligand-dependent suppression of Cas12a genome editing, with InDel frequencies dropping by about 50 % upon COR addition, while positive and negative controls were again unaffected by the ligand (Figure 4F and G). Of note, the overall potency of CASANOVA-VA1 in presence of the drug was significantly lower than that of wild-type AcrVA1. Nevertheless, our data indicate that both, the CASASNOVA concept for optogenetic Acr inhibition as well as our novel CASANDRA concept for ligand-dependent Acr activation can, in principle, be applied to Acrs from different CRISPR-Cas clades.

## DISCUSSION

The ability to control and modulate CRISPR effector activity in living cells through exogenous triggers remains a key challenge in the field. Despite significant advancements, there is still a lack of modular and flexible tools capable of regulating the wide array of CRISPR effectors currently used across diverse applications in the life sciences. Most available control strategies are tailored to specific CRISPR-Cas orthologs or limited to particular use cases. In this study, we introduced a straightforward, Acr-based platform for opto- and chemogenetic regulation of diverse type II and type V CRISPR effectors, leveraging the remarkable broad-spectrum activity of certain Acrs.

Our domain insertion screen identified several regions in AcrIIA5 that well-tolerate LOV2 domain fusion, indicating high structural flexibility of AcrIIA5 in accommodating an additional protein domain. Apart from the fusion sites directly at or close to the N- and C-termini of AcrIIA5, domain insertion tolerance was highest at the position preceding N77, which lies directly within the second intrinsically disordered region (IDR; residues ∼65-79) of the AcrIIA5 protein (Supplementary Figure S3). This observation well aligns with a point mutation scanning dataset recently reported by us, in which the this internal IDR was identified as being a particularly mutation tolerant region (38).

In stark contrast, positions tested for LOV2 insertion within the core part of the first, N-terminal IDR of AcrIIA5 did not tolerate LOV2 fusion at all (Supplementary Figure S3, residues 14-20). This suggests that domain fusion at these sites likely resulted in steric clashes, either within AcrIIA5 itself or with Cas9. Given the structural flexibility of the IDR and the hypothesis that the N-terminal IDR in AcrIIA5 functions as a Cas9 tether (37,38), the latter explanation seems plausible. This interpretation is further supported by the observation that surface regions of AcrIIA5 adjacent to the N-terminal IDR, i.e., “facing the IDR”, also were rather intolerant to LOV2 fusion. This aligns with the proposed IDR-mediated tethering mechanism of AcrIIA5: When the IDR binds to Cas9, the surface sites of AcrIIA5 adjacent to the N-terminal IDR would likely be positioned close to the Cas9 protein surface. Domain insertion into these IDR-adjacent sites of the AcrIIA5 structure would thus result in steric clashes with Cas9, explaining their inability to tolerate domain fusion. While still rather speculative at this point, these observations are consistent with previous mechanistic evidence (37,38) and provide further support for the proposed IDR-mediated tethering mechanism in AcrIIA5 function.

The CASANOVA-A5 and CASANDRA-A5 variants generally demonstrated robust regulation of various Cas9 orthologs across different human cell lines. Throughout our experiments, however, we observed an intrinsic balance between the ability of different CASANOVA-A5 and CASANDRA-A5 variants to effectively inhibit Cas9 in the off-state and release its activity in the on-state (e.g., compare the D41 and N77 variants in Figures 1 and 2). This trade-off may partially stem from the inherent characteristics of the used, allosteric control mechanism itself. For the LOV2, for instance, it is known that this photosensor exists in an equilibrium between the dark and lit states, with photoexcitation strongly shifting the balance toward the lit state (59,60). Such equilibrium behavior likely contributes to the observed baseline leakiness and dynamic range of control in both, CASANOVA and CASANDRA.

A second, important factor influencing performance of our switchable Acrs is the co-expression of CASANOVA/CASANDRA with the cognate Cas effector and RNA guide, which requires an optimal ratio of the different components for maximal efficacy. We observed, for instance, cell type-specific differences in regulation efficiency among HEK293T, HCT116, and HeLa cells, likely reflecting variations in transfection efficiency and overall expression strength. Importantly, transient transfection inherently produces a heterogeneous population of cells with varying expression levels of each component. Alternative delivery strategies, such as AAV transduction, could achieve more uniform co-expression of Cas9/Cas12, sgRNA/crRNA, and CASANOVA/CASANDRA across cells. Such approaches have previously shown promise for enhancing performance, as exemplified by our earlier observations with AAV delivery of CASANOVA-A4 (14), but due to the AAV cargo size limit, may require multiple AAV vectors.

In this context, we were excited to see that it is generally possible to hijack the pol-II activity of pol-III promoters for co-expression of multiple proteins, i.e. CASANOVA and *Sau*Cas9, alongside the guide RNA (Supplementary Figure S5). While conceptually very interesting, the real-world potential of this strategy in context of AAV delivery will require detailed exploration through future work. This is particularly true since current AAV CRISPR vectors usually rely – for good reasons - on very potent pol-II promoters for Cas effector expression, such as CMV or EF1α promoters. These are sometimes even topped-up with additional transcription-enhancing elements (61), since high Cas effector expression appears to be critical for effective genome editing with AAVs in cell culture and *in vivo*. Nevertheless, the fact that we could “squeeze” *Sau*Cas9, an Acr-LOV2 hybrid and a sgRNA into one transgene with a size below the AAV cargo limit is an interesting starting point for further research towards rationally engineering or, through direct evolution, creating new, potent hybrid pol-II/pol-III promoters. These could then be used to effectively co-express CRISPR components and additional control systems like our CASANOVA/CASANDRA system from single AAV vectors.

On top of controlling diverse Cas9 orthologs with exogenous triggers, our work also provides the first proof-of-concept for Cas12a opto- and chemogenetic control with switchable AcrVA1, expanding our CASANOVA/CASANDRA concept to the type V CRISPR effector clade. Of note, for AcrVA1, we have thus far only tested a few insertion site candidates within a region identified through inspection of the AcrVA1/Cas12a structure. While insertion of the LOV2 and cpGR2 domain into this region readily yielded switchable AcrVA1 variants, their performance lagged behind that obtained by the more extensive screening in AcrIIA5. Therefore, it would be interesting to explore additional, surface-exposed sites in AcrVA1 for domain insertion tolerance, e.g. by systematic insertion screening (62), and/or extend this approach to other type V Acrs.

The cpReceptor domains developed here hold significant promise for engineering protein switches in synthetic biology, potentially compatible with other soluble receptors reported as split-variants (45). Notably, cpReceptor domains can replace the LOV2 domain in previously identified allosteric sites (Figure 4), indicating broad potential for cpReceptor-based chemical regulation beyond Acrs.

Notably, unlike LOV2, which typically generates light-inactivated effectors due to local disorder introduced upon helix unfolding in response to light (63) (albeit exceptions exist (62,64)), the cpReceptors function inversely: In their ligand-unbound state, cpReceptors are flexible, while ligand binding induces a compact conformation with proximate receptor termini. This structural behavior makes our engineered cpReceptors particularly well-suited for engineering ligand-activated protein switches.

In summary, the extension of CASANOVA to broad-spectrum CRISPR-Cas9 inhibitors and type V CRISPR effectors, along with the complementary development of CASANDRA for chemical Acr regulation, provides a versatile toolbox for opto- and chemogenetic control of diverse CRISPR systems. More broadly, our work paves the path towards modular engineering of allosteric effector proteins controlled with various inputs.

## DATA AVAILABILITY

DNA and Protein sequences (Genbank files) of CASANOVA/CASANDRA constructs are provided as Supplementary Data 1, and relevant plasmids encoding these will be made available via Addgene. Otherwise, all relevant data is contained directly within the manuscript.

## ACKNOWLEDGEMENTS

We thank all members of the Niopek lab for helpful discussions. We thank Klara Eisenhauer, Technical University of Darmstadt, for supporting initial experiments performed by C.G.. *Author contributions:* D.N. conceived the study and refined it with J.M., L.B., F.B., S.A. and C.G.. L.B., F.B., S.A., C.G., J.M and D.N. designed the experiments. L.B., F.B., S.A., A-S.K and K.S. performed experiments. C.G. cloned the cpTRβ and cpERβ constructs and performed initial experiments. L.B., J.M. and M.J. analyzed data. B.W. performed protein structure predictions and color-mapping. All authors contributed to data interpretation. D.N. directed the work and secured funding. L.B. and D.N. wrote the manuscript. All authors approved the final manuscript.

## SUPPLEMENTARY DATA LEGENDS

**Suppelementary Data 1:** Genbank files with annotated DNA sequences of CASANOVA and CASANDRA constructs (refer to Supplementary Table T1).

