## Supplementary Information for "A Versatile Anti-CRISPR Platform for Opto- and Chemogenetic Control of CRISPR-Cas9 and Cas12 across a Wide Range of Orthologs"

**SUPPLEMENTARY FIGURES**

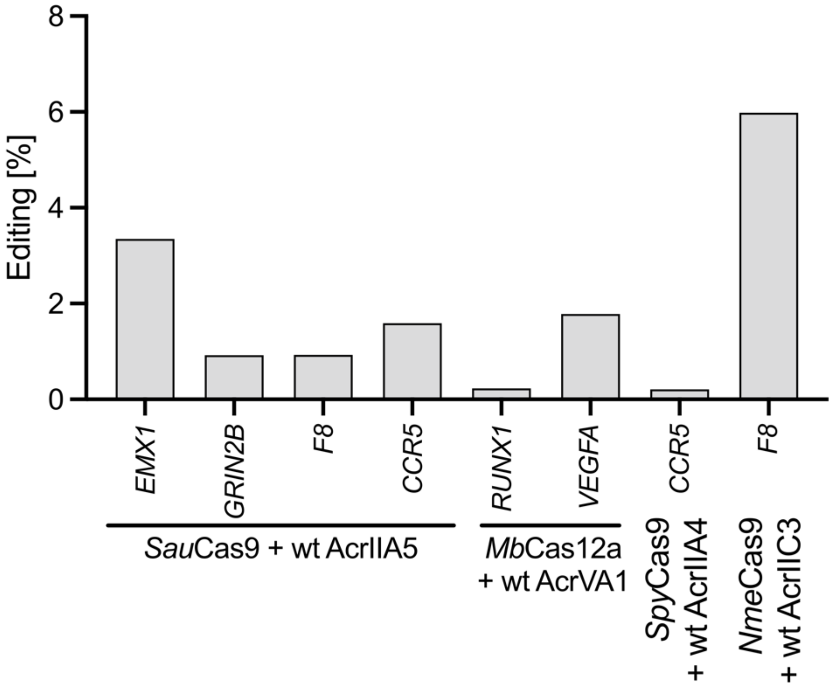


**Supplementary Figure S1: Assessing leakiness of Cas effector inhibition with wild-type Acrs.** HEK293T cells were co-transfected with plasmids encoding (i) the indicated CRISPR-Cas and a sgRNA/crRNA targeting the indicated locus and (ii) the indicated wt Acr. Cells were incubated for 72 h, followed by amplification of the target locus with unique double-indexed primers and subsequent NGS analysis of InDels at the targeted locus. Data corresponds to a single experiment. The specific Cas effector:target site:Acr pairs are as follows: *Sau*Cas9:*EMX1*:wt AcrIIA5; *Sau*Cas9:*GRIN2B*:wt AcrIIA5; *Sau*Cas9:*F8*:wt AcrIIA5; *Sau*Cas9:*CCR5*:wt AcrIIA5; *Mb*Cas12a:*RUNX1*:wt AcrVA1; *Mb*Cas12a:*VEGFA*:wt AcrVA1; *Spy*Cas9:*CCR5*:wt AcrIIA4; *Nme*Cas9:*F8*:wt AcrIIC3. Wt, wild-type.

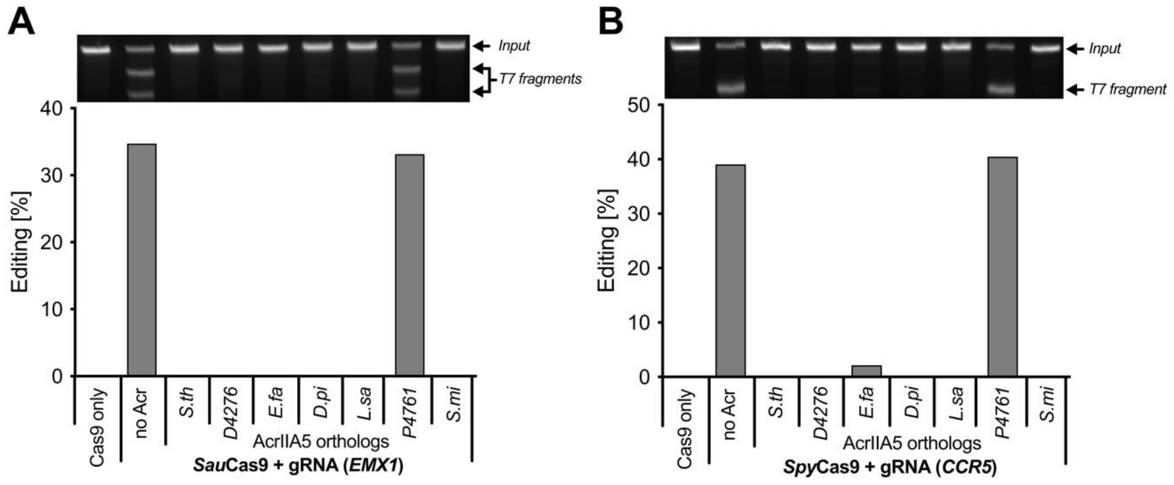


**Supplementary Figure S2. Inhibition potency of different AcrIIA5 orthologs.** (A, B) HEK293T cells were co-transfected with plasmids encoding (i) *Sau*Cas9 and a sgRNA targeting the *EMX1* locus (A) or *Spy*Cas9 and a sgRNA targeting the *CCR5* locus (B) and (ii) various wild-type AcrIIA5 orthologs derived from bacteriophages (identified by their host bacterial strains). Cells were incubated for 72 h post transfection, followed by T7E1 analysis of InDels at the targeted locus. Data corresponds to a single experiment. AcrIIA5 orthologs originate from the following bacteriophage host stains: *Streptococcus thermophilus (S.th)*; *Streptococcus pyogenes* strain *D4276*; *Enterococcus faecalis (E.fa)*; *Dolosigranulum pigrum (D.pi)*; *Lactobacillus sakei (L.sa)*; *Streptococcus pneumoniae* strain *P4761* and *Streptococcus mitis (S.mi)*.

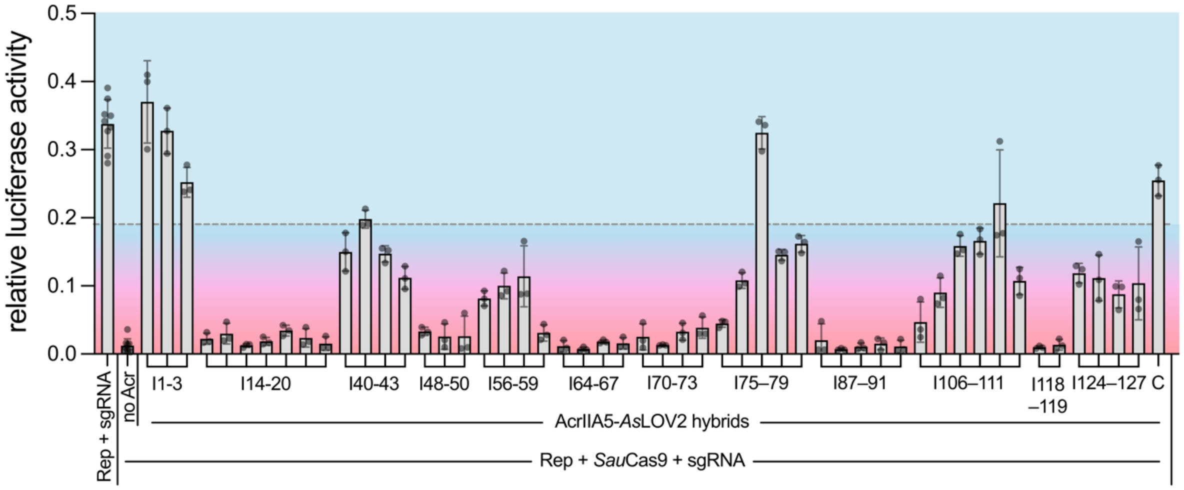


**Supplementary Figure S3: *As*LOV2 domain insertion screen in AcrIIA5 identifies sites tolerating domain fusion.** HEK293T cells were co-transfected with plasmids encoding (i) *Sau*Cas9 and a sgRNA targeting a firefly luciferase, (ii) the indicated AcrIIA5-*As*LOV2 hybrids (residues following the LOV2 insertion site are indicated) (iii) a dual luciferase reporter (Rep) co-expressing the firefly and *Renilla* luciferase. Cells were incubated for 48 h in the dark, followed by luciferase assay. Firefly photon counts were normalized to *Renilla* photon counts. Bars indicate means, grey dots individual data points, and error bars the SD from n = 3 technical replicates (sample groups) or n = 9 technical replicates (controls – note that controls were always included on well-plates, each carrying different sample groups).

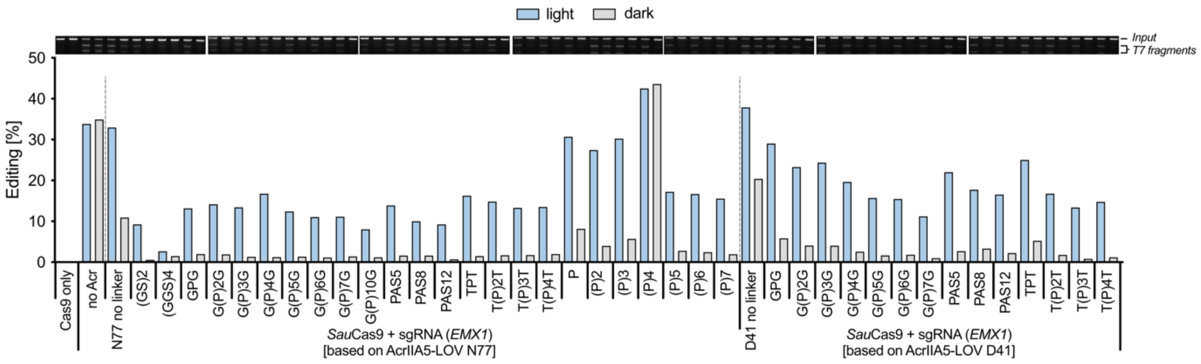


**Supplementary Figure S4**: **Symmetric linkers appended to *As*LOV2:Acr junction sites in CASANOVA-A5 variants improve switch performance.** HEK293T cells were co-transfected with plasmids encoding the indicated components and illuminated for 72 h or kept in the dark, followed by T7E1 analysis of InDels at the targeted *EMX1* locus. AcrIIA5-*As*LOV2 hybrids are indicated by the appended symmetric GS-rich flexible, semi-rigid (e.g. PAS and TPT) or rigid proline-based linkers respectively. Data corresponds to a single experiment.

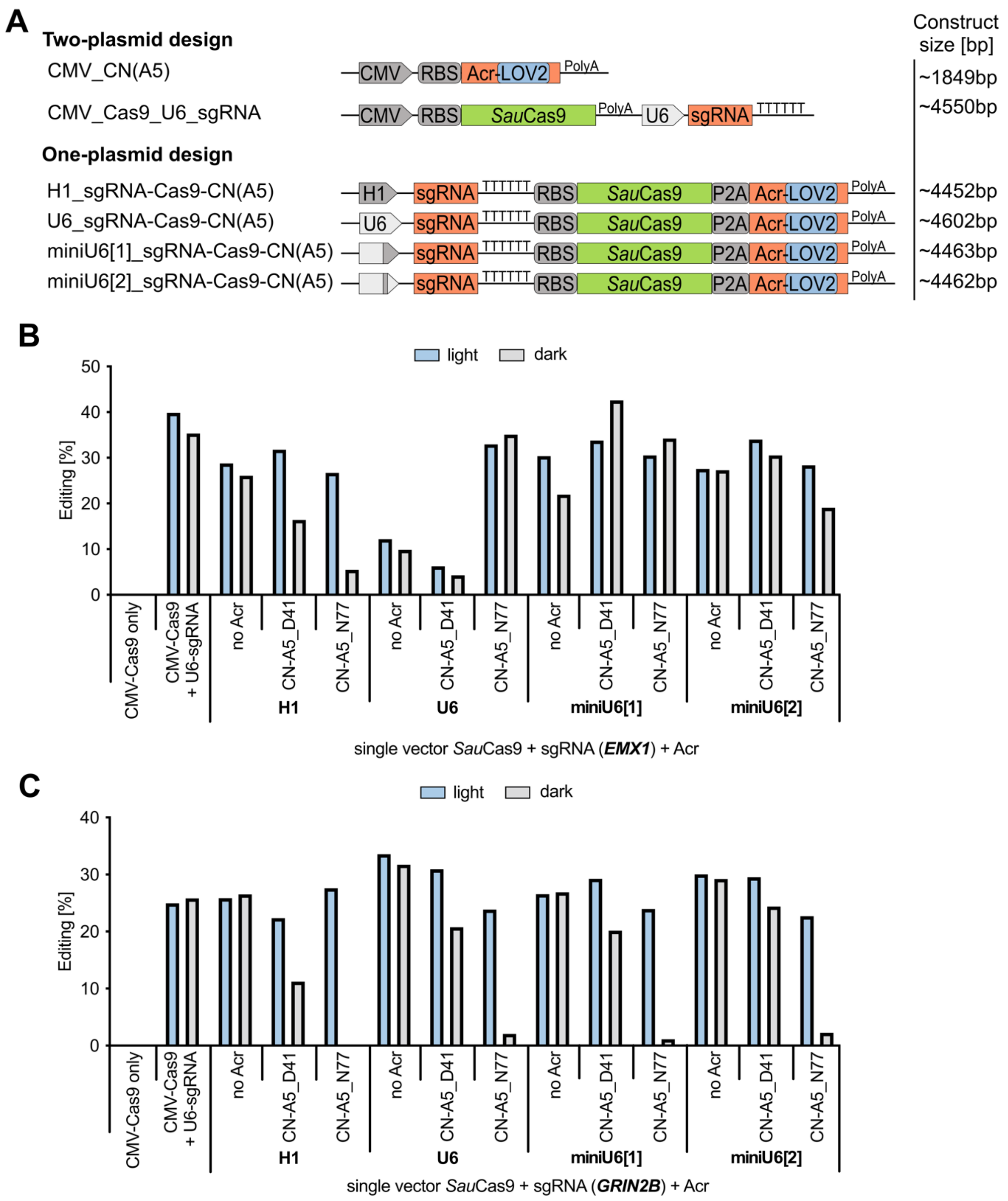


**Supplementary Figure S5: Expression of all components of the CASANOVA-A5 system from a single pol-III promoter.** (A) Schematic of two-plasmid and one-plasmid vector designs and corresponding base-pair length of the encoded transgene(s). (B, C) HEK293T cells were transfected with one-plasmid designs depicted in A and illuminated for 72 h or kept in the dark, followed by T7E1 analysis of InDels at the targeted *EMX1* (B) or *GRIN2B* (C) locus. Data corresponds to a single experiment. MiniU6 promoters were constructed as either: (i) a minimal U6 promoter fused to the 3' 29 bp of the H1 promoter (miniU6[1]), or (ii) a minimal U6 promoter with its TATA Box replaced by that of the H1 promoter (miniU6[2]). See Supplementary Table S4 for promoter sequences. CN: CASANOVA. D41/N77 identify the positions prior to which LOV2 was inserted into AcrIIA5 in the one-plasmid design.

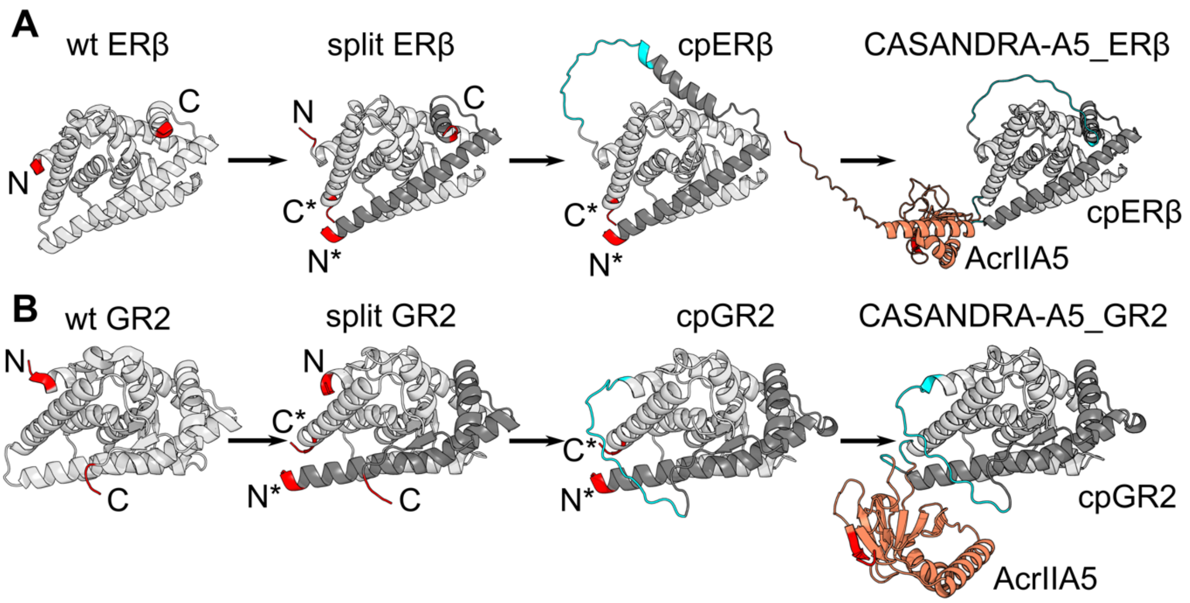


**Supplementary Figure S6: Design of CASANDRA-A5 variants based on ERβ and GR2 receptor domains.** (A, B) In analogy to the schematic in Figure 2A, cpReceptor engineering stages are shown for the human estrogen receptor beta domain (ERβ; PDB ID: 3ols; first three structures from the left) (A) and the glucocorticoid receptor 2 domain (GR2; PDB ID: 4e2j; first three structures from the left) (B). The structure on the right shows an AlphaFold3 prediction of the fusion between cpERβ and AcrIIA5 (D41 variant) (A) or cpGR2 and AcrIIA5 (N77 variant) (B); red: termini; blue: linker residues.

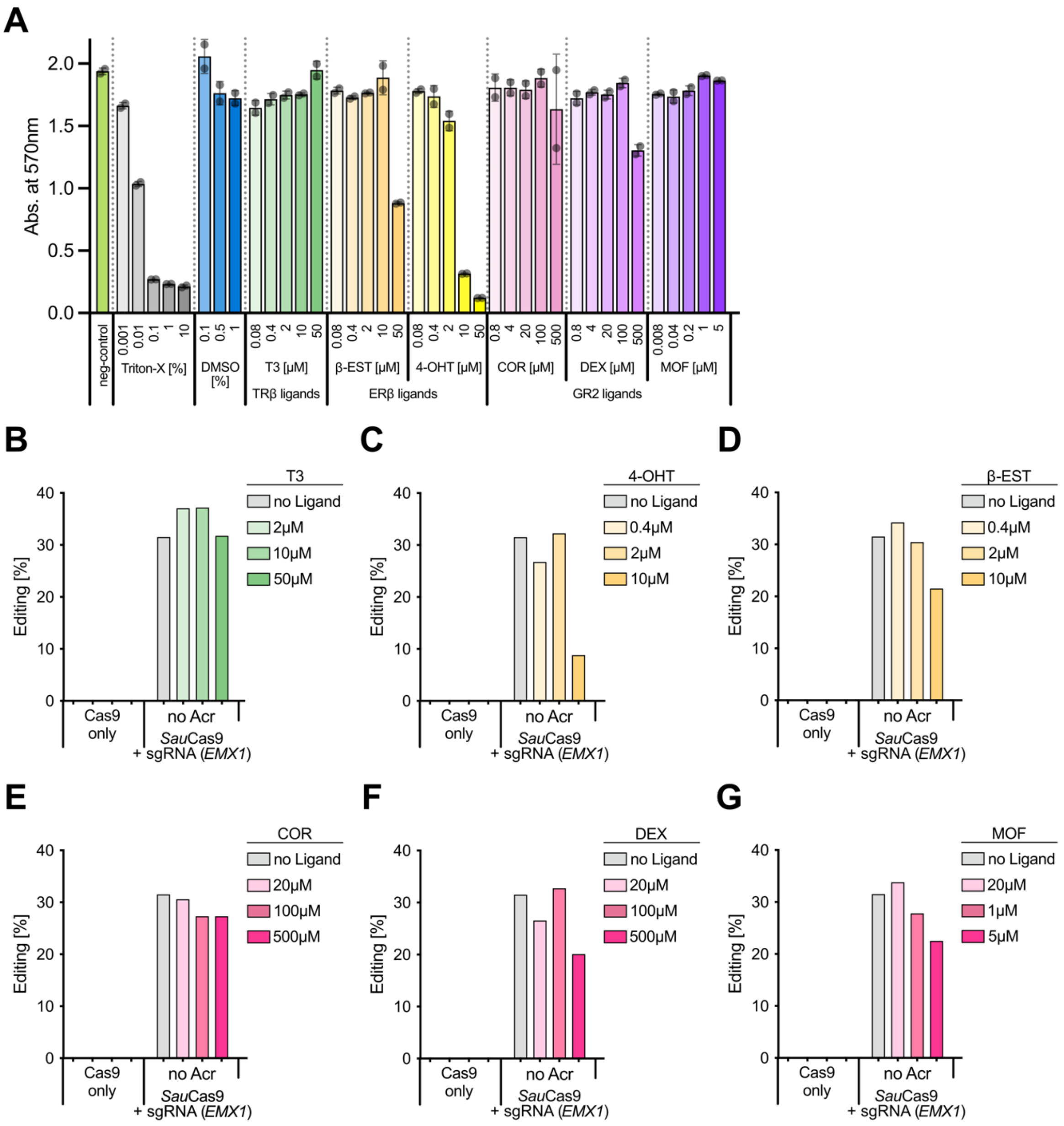


**Supplementary Figure S7: Assessment of mammalian cell toxicity of the used drugs and their effect on CRISPR genome editing.** (A) HEK293T cells were treated with the indicated drugs at indicated concentrations and incubated for 72 h, followed by MTT assay to assess cell viability. Bars indicate means, grey dots individual data points, and error bars the SD from n = 3 different replicates (wells). The specific ligands are as follows: triiodothyronine (T3); 4-hydroxytamoxifen (4-OHT); β-estradiol (β-EST); cortisol (COR); dexamethasone (DEX) and mometasone furoate (MOF). Various concentrations of Triton X-100 were used as control for cell death and untreated cells as control for maximum viability. (B-G) HEK293T cells were co-transfected with the indicated components and treated with T3 (B); 4-OHT (C); β-EST (D); COR (E); DEX (F) or MOF (G) starting 2 h later. 72 h post transfection, cells were lysed, followed by NGS analysis of InDels at the targeted *EMX1* locus. Data corresponds to a single experiment.

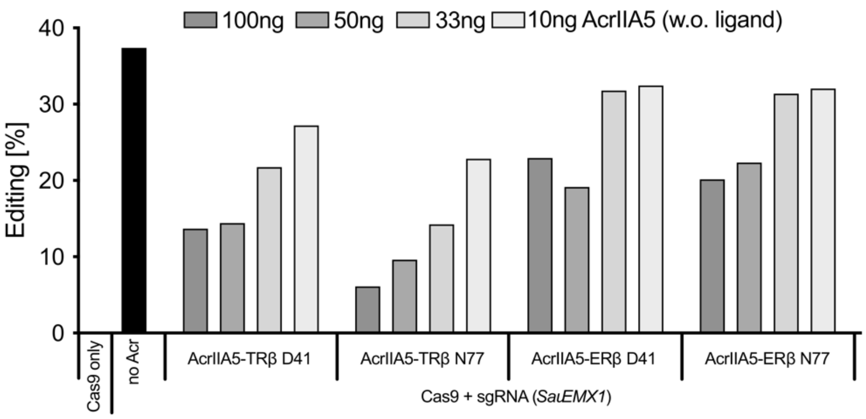


**Supplementary Figure S8: Plasmid titration of AcrIIA5-cpReceptor variants and resulting effects on *Sau*Cas9-mediated genome editing.** HEK293T cells were co-transfected with plasmids encoding (i) *Sau*Cas9 and a sgRNA targeting the *EMX1* locus and (ii) the indicated AcrIIA5-cpReceptor variants using the indicated plasmid amount per well. Cells were incubated for 72 h post transfection, followed by T7E1 analysis of InDels at the *EMX1* locus. Data corresponds to a single experiment.

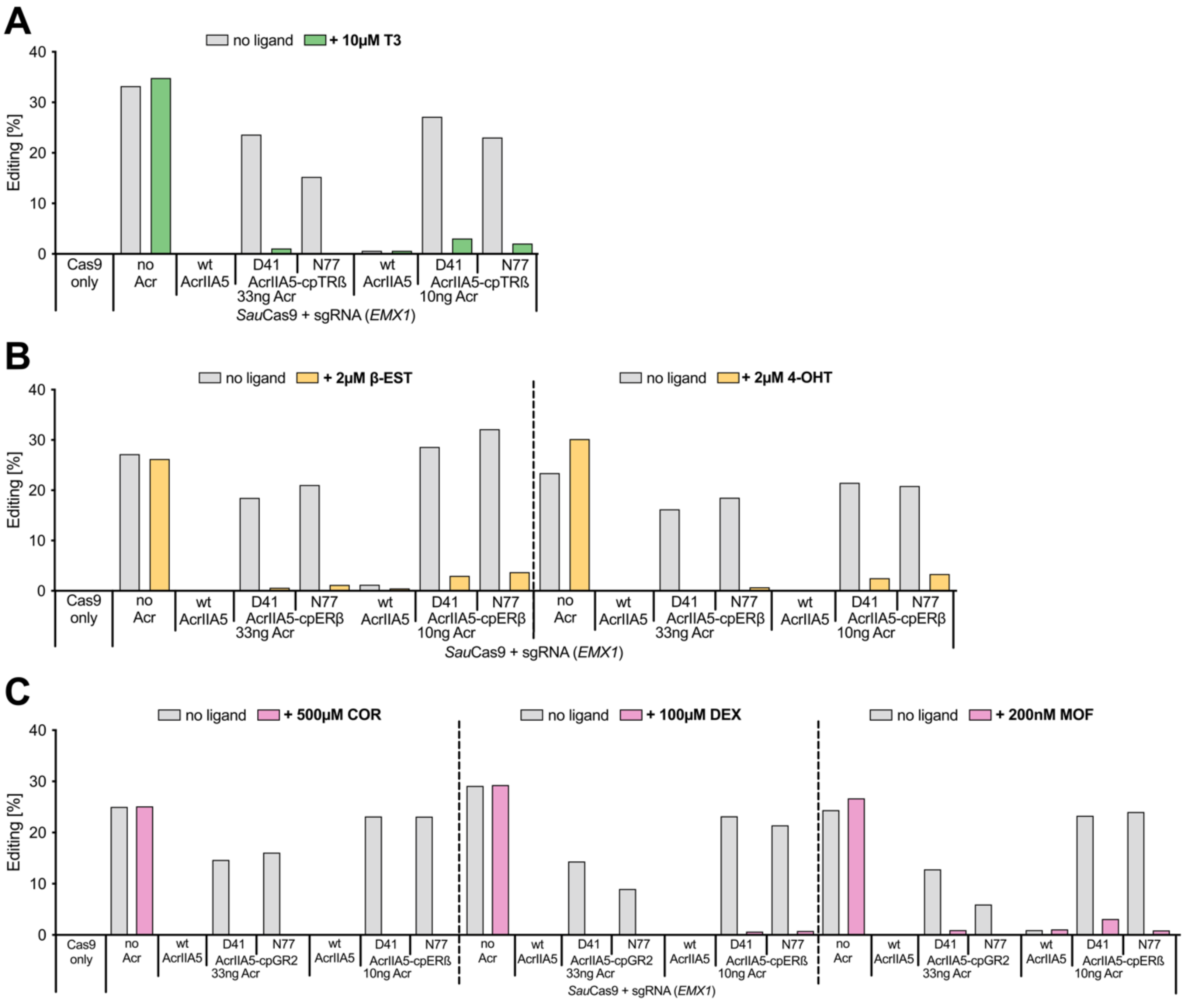


**Supplementary Figure S9: Impact of CASANDRA-A5 plasmid amounts used for transfection on *Sau*Cas9-mediated editing and ligand-dependent control.** (A-C) HEK293T cells were co-transfected with plasmids encoding (i) *Sau*Cas9 and a sgRNA targeting the *EMX1* locus and (ii) indicated AcrIIA5-cpReceptor variants using 33 ng or 10 ng of plasmid for transfection. Cells were treated with the indicated drugs starting 2 h later. 72 h post transfection, cells were lysed, followed by T7E1 analysis of InDels at the targeted locus. Data corresponds to a single experiment. The specific cpReceptor:ligand pairs are as follows. (A) cpTRβ:T3. (B) cpERβ: β-EST and cpERβ:4-OHT. (C) cpGR2:COR, cpGR2:DEX and cpGR2:MOF.

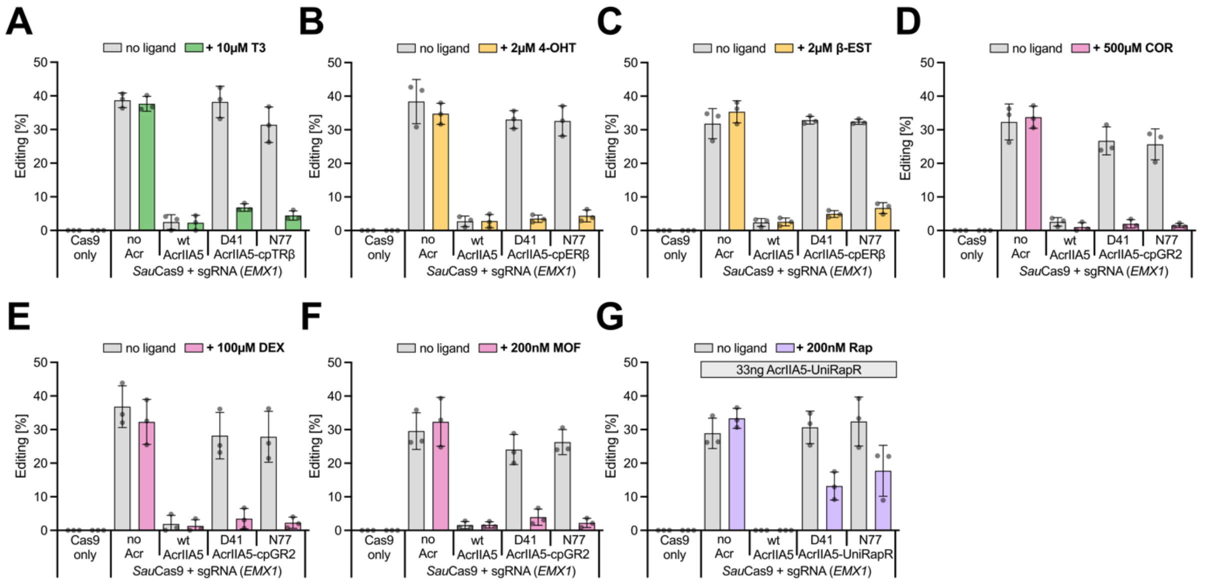


**Supplementary Figure S10: Performance of CASANDRA-A5 and comparison with a variant based on the UniRapR domain.** (A-G) HEK293T cells were co-transfected with plasmids encoding (i) *Sau*Cas9 and a sgRNA targeting the *EMX1* locus and (ii) the indicated AcrIIA5-cpReceptor using 10 ng (A-F) or 33 ng (G) of plasmid for transfection per well. Cells were treated with the indicated drugs starting 2 h later. 72 h post transfection, cells were lysed, followed by T7E1 analysis of InDels at the targeted locus. Bars indicate means, grey dots individual data points, and error bars the SD from n = 3 independent experiments. The specific cpReceptor:ligand pairs are as follows. (A) cpTRβ:T3. (B) cpERβ:4-OHT. (C) cpERβ: β-EST. (D) cpGR2:COR. (E) cpGR2:DEX. (F) cpGR2:MOF and (G) UniRapR:Rapamycin (Rap).

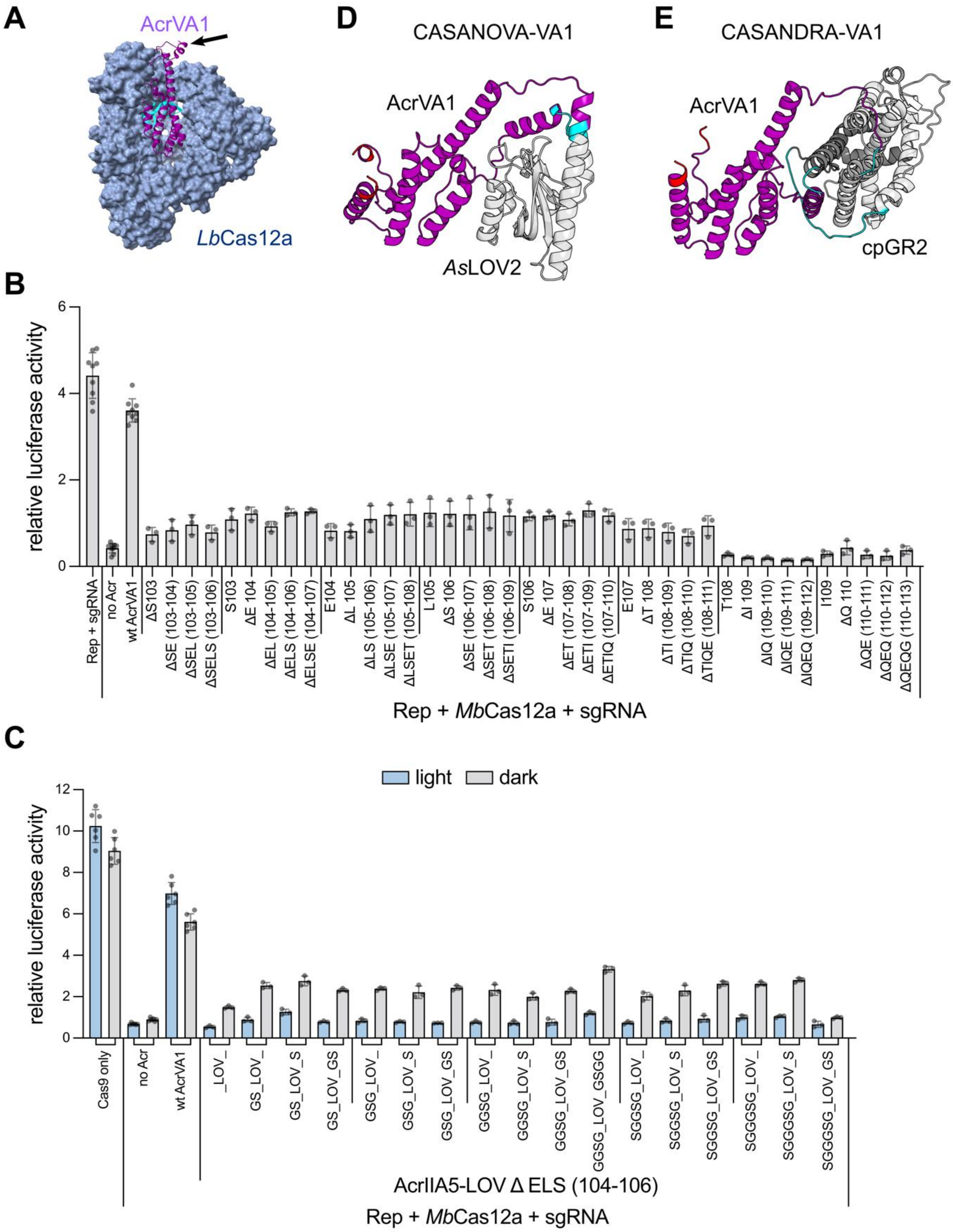


**Supplementary Figure S11: LOV2 domain insertion yields photosensitive variants of AcrVA1, a Cas12a inhibitor.** (A) Schematic of *Lb*Cas12a (blue) in complex with AcrVA1 (purple) (PDB ID: 6nmd). The surface sites in AcrVA1 assessed for LOV2 domain insertion is indicated with an arrow. Note that we show the *Lb*Cas12a-AcrVA1 complex here, since for *Mb*Cas12a, there is no structure available in complex with this Acr. (B) HEK293T cells were co-transfected with plasmids encoding (i) *Mb*Cas12a and a sgRNA targeting a firefly luciferase, (ii) the AcrVA1-*As*LOV2 hybrids indicated by insertion positions following the insertion site and their cognate deletions around the insertion site, (iii) a luciferase reporter expressing a firefly luciferase and (iv) a vector expressing a *Renilla*

luciferase. Cells were incubated for 48 h in the dark, followed by luciferase assay. Firefly photon counts were normalized to *Renilla* photon counts. Bars indicate means, grey dots individual data points, and error bars the SD from n = 3 technical replicates (sample groups) or n = 9 technical replicates (controls – note that controls were always included on well-plates, each carrying different sample groups). (C) HEK293T cells were co-transfected with plasmids encoding (i) *Mb*Cas12a and a sgRNA targeting a firefly luciferase, (ii) the AcrVA1-*As*LOV2 ΔELS (104-106) variant with indicated linkers connecting the Acr:*As*LOV2 junction sites, (iii) a luciferase reporter expressing a firefly luciferase and (iv) a vector expressing a *Renilla* luciferase. Cells were illuminated for 48 h or kept in the dark, followed by luciferase assay analysis. Firefly photon counts were normalized to *Renilla* photon counts. Bars indicate means, grey dots individual data points, and error bars the SD from n = 3 technical replicates (sample groups) or n = 9 technical replicates (controls – note that controls were always included on well-plates, each carrying different sample groups).

### SUPPLEMENTARY TABLES

**Supplementary Table T1: List of most important plasmids used and created in this study.** Numbers indicate plasmids created in this work and with corresponding genbank files available as Supplementary Data 1. Grey labels indicate vectors not included in Supplementary Data 1.

| .gb # | Vector description | Vector name | Source |
|---|---|---|---|
| | Firefly and *Renilla* luciferase reporter | pAAVpsi2 | Bubeck 2018 |
| 1 | Firefly luciferase only reporter | pSi-AAV w/o-Renilla | This study |
| | *Renilla* only reporter | pRL-TK | Promega |
| | CMV-*Sau*Cas9-U6-SauCas9 sgRNA scaffold | pX601-AAV-CMV::NLS-SaCas9-NLS-3xHA-bGHpA;U6::BsaI-sgRNA | Addgene #61591 |
| 2 | *Sau*Cas9 + sgRNA targeting FFLuc | SaCas9_U6_gRNA(FF1) | This study |
| 3 | SauCas9 + sgRNA targeting *EMX1* | SaCas9_U6_gRNA(EMX1) | This study |
| 4 | SauCas9 + sgRNA targeting *GRIN2B* | SaCas9_U6_gRNA(GRIN2B) | This study |
| | *Nme*Cas9 + sgRNA targeting *F8* | pEJS654 All-in-One AAV-sgRNA_F8-hNmeCas9 | Hoffmann 2021 |
| | *Spy*Cas9 expression vector | pcDNA3.1 3xFlag-NLS-SpCas9-NLS | Bubeck 2018 |
| | sgRNA targeting CCR5 with *Spy*Cas9 scaffold | U6-gRNA CCR5 | Bubeck 2018 |
| | *Mb*Cas12a expression vector | pCAG-hMb3Cas12a-NLS(nucleoplasmin)-3xHA (RTW2500) | Addgene #115142 |
| 5 | *Mb*Cas12a + crRNA targeting firefly luciferase | MbCas12a U6_crRNA (FFLuc) | This study |
| 6 | *Mb*Cas12a + crRNA targeting *RUNX1* | MbCas12a U6_crRNA (RUNX1) | This study |

| | | | |
|---|---|---|---|
| 7 | *Mb*Cas12a + crRNA targeting *VEGFA* | MbCas12a U6_crRNA (VEGFA) | This study |
| 8 | Wild-type AcrIIA5 from *Streptococcus thermophilus* | CMV_AcrIIA5 | This study |
| | Wild-type AcrIIA4 | pcDNA3.1 SV40NLS-GGS-AcrIIA4 | Bubeck 2018 |
| | Wild-type AcrIIC3 | pcDNA3.1_AcrIIC3-miRNA-scaffold-BsmBI | Hoffmann 2021 |
| 9 | Wild-type AcrVA1 | CMV_AcrVA1 | This study |
| 10 | AcrIIA5-*As*LOV2 insertion preceding position D41 | CMV_AcrIIA5_LOV_InsD41 | This study |
| 11 | AcrIIA5-*As*LOV2 insertion preceding position N77 | CMV_AcrIIA5_LOV_InsN77 | This study |
| 12 | AcrIIA5-*As*LOV2; exemplary vector with altered linkers flanking LOV2 | AcrIIA5-AsLOV2(D41)-GPG | This study |
| 13 | AcrIIA5-cpTRβ insertion D41 | pCMV-AcrIIA5_cpTRb(GGS)5_D41 | This study |
| 14 | AcrIIA5-cpTRβ insertion N77 | pCMV-AcrIIA5_cpTRb(GGS)5_N77 | This study |
| 15 | AcrIIA5-ERβ insertion D41 | pCMV-AcrIIA5_cpERb(GGS)5_I41 | This study |
| 16 | AcrIIA5-ERβ insertion N77 | pCMV-AcrIIA5_cpERb(GGS)5_I77 | This study |
| 17 | AcrIIA5-GR2 insertion D41 | pCMV_AcrIIA5_cpGR2(GGS5)_I41 | This study |
| 18 | AcrIIA5-GR2 insertion N77 | pCMV_AcrIIA5_cpGR2(GGS5)_I77 | This study |
| 19 | AcrVA1-LOV ΔELS (104-106) | AcrVA1_*As*LOV2-S103ΔELS | This study |
| 20 | AcrVA1-*As*LOV; exemplary vector with altered linkers flanking LOV2 | AcrVA1_(SGGSG)LOV2(GS)- S103ΔELS | This study |

| | | | |
|---|---|---|---|
| 21 | AcrVA1-GR2 insertion with GPG Linker | AcrVA1_cpGR2(GGS5)-S103ΔELS | This study |
| 22 | AcrVA1-GR2 insertion with (GGS)2 Linker | AcrVA1_cpGR2(GGS5)-S103ΔELS_(GGS)2-(GGS)2 | This study |
| 23 | AcrIIA4-GR2 insertion | pCMV_AcrIIA4_cpGR2(GGS5)_ΔN64/Q65/E66 | This study |
| 24 | AcrIIC3-GR2 insertion | pCMV_AcrIIC3_cpGR2(GGS5)_F59 | This study |
| 25 | H1-*EMX1* with *Sau*Cas9 (pol-III single plasmid design; see Supplementary Figure S5) | H1_sgRNA(EMX1)_SauCas9 | This study |
| 26 | H1-*GRIN2B* with SauCas9 (pol-III single plasmid design; see Supplementary Figure S5) | H1_sgRNA(GRIN2B)_SauCas9 | This study |
| 27 | H1-*EMX1 Sau*Cas9-P2A-CASANOVA-A5(D41) (pol-III single plasmid design; see Supplementary Figure S5) | H1_sgRNA(EMX1)_SauCas9_AcrIIA5-AsLOV2(D41) | This study |
| 28 | H1-*EMX1 Sau*Cas9-P2A-CASANOVA-A5(N77) (po-III single plasmid design; see Supplementary Figure S5) | H1_sgRNA(EMX1)_SauCas9_AcrIIA5-AsLOV2(N77) | This study |
| 29 | H1-*GRIN2B Sau*Cas9-P2A-CASANOVA-A5(D41) (pol-III single plasmid design; see Supplementary Figure S5) | H1_sgRNA(GRIN2B)_SauCas9_AcrIIA5-AsLOV2(D41) | This study |
| 30 | H1- *GRIN2B Sau*Cas9-P2A-CASANOVA-A5(N77) (pol-III single plasmid design; see Supplementary Figure S5) | H1_sgRNA(GRIN2B)_SauCas9_AcrIIA5-AsLOV2(N77) | This study |
| 31 | U6-*EMX1 Sau*Cas9 (pol-III single plasmid design; see Supplementary Figure S5) | U6_sgRNA(EMX1)_SauCas9 | This study |

| | | | |
|---|---|---|---|
| 32 | miniU6[1]-*EMX1 Sau*Cas9 (pol-III single plasmid design; see Supplementary Figure S5) | miniU6[1]_sgRNA(EMX1)_SauCas9 | This study |
| 33 | miniU6[2]-*EMX1 Sau*Cas9 (pol-III single plasmid design; see Supplementary Figure S5) | miniU6[2]_sgRNA(EMX1)_SauCas9 | This study |
| | DNA stuffer plasmid | pBluescript sk- | Invitrogen |

**Supplementary Table T2: cpReceptor sequences used in CASANDRA variants**

cpTRβ, cpERβ and GR2 were tested in this study. cpPPAR*γ* is presented as exemplary additional cpReceptor candidate that follows the analogous design, but was not tested in this study.

| | | |
|---|---|---|
| cpTRβ | cTRβ (214-261)-(GGS)5-nTRβ (1-213) | HVTHFWPKLLMKVTDLRMIGACHASRFLHMKVECPTELFPPLFLEVFEGGGSGGSGGSGGSGGSGHMEELQKSIGHKPEPTDEEWELIKTVTEAHVATNAQGSHWKQKRKFLPEDIGQAPIVNAPEGGKVDLEAFSHFTKIITPAITRVVDFAKKLPMFCELPCEDQIILLKGCCMEIMSLRAAVRYDPESETLTLNGEMAVTRGQLKNGGLGVVSDAIFDLGMSLSSFNLDDTEVALLQAVLLMSSDRPGLACVERIEKYQDSFLLAFEHYINYRKH |
| cpERβ | cERβ (187-241)-(GGS)5-nERβ (1-186) | SSQQQSMRLANLLMLLSHVRHASNKGMEHLLNMKCKNVVPVYDLLLEMLNAHVLRGGGSGGSGGSGGSGGSGDALSPEQLVLTLLEAEPPHVLISRPSAPFTEASMMMSLTKLADKELVHMISWAKKIPGFVELSLFDQVRLLESCWMEVLMMGLMWRSIDHPGKLIFAPDLVLDRDEGKCVEGILEIFDMLLATTSRFRELKLQHKEYLCVKAMILLNSSMYPLVTATQDADSSRKLAHLLNAVTDALVWVIAKSGI |
| cpGR2 | cGR2(176-247)-(GGS)5-nGR2(1-175) | NSSQNWQRFYQLTKLLDSMHEMVGGLLQFCFYTFVNKSLSVEFPEMLAEIISNQLPKFNAGSVKPLLFHQKGGGSGGSGGSGGSGGSGLISLLEVIEPEVLYSGYDSTLPDTSTRLMSTLNRLGGRQVVSAVKWAKALPGFRNLHLDDQMTLLQYSWMSLMAFSLGWRSYKQSNGNMLCFAPDLVINEERMQLPYMYDQCQQMLKISSEFVRLQVSYDEYLCMKVLLLLSTVPKDGLKSQAVFDEIRMTYIKELGKAIVKREG |
| cpPPAR*γ* | cPPAR*γ*(231-278)-(GGS)5-nPPAR*γ*(1-230) | QLFAKLLQKMTDLRQIVTEHVQLLQVIKKTETDMSLHPLLQEIYKDLYGGGSGGSGGSGGSGGSGGSHMLNPESADLRALAKHLYDSYIKSFPLTKAKARAILTGKTTDKSPFVIYDMNSLMMGEDKIKFKHITPLQEQSKEVAIRIFQGCQFRSVEAVQEITEYAKSIPGFVNLDLNDQVTLLKYGVHEIIYTMLASLMNKDGVLISEGQGFMTREFLKSLRKPFGDFMEPKFEFAVKFNALELDDSDLAIFIAVIILSGDRPGLLNVKPIEDIQDNLLQALELQLKLNHPESS |

**Supplementary Table T3: sgRNA/crRNA genomic target sequences**

Sequences are in 5’ to 3’ orientation; the sequence marked in bold represents the respective PAM

| | | |
|---|---|---|
| *Sau*Cas9 | *Luciferase* | cactggcatgaagaactgcaga**gagt** |
| | *EMX1* | ggcctccccaaagcctggccagg**gagt** |
| | *GRIN2B* | gagtaggctggtagatggagtt**gggt** |
| *Spy*Cas9 | *CCR5* | tgacatcaattattatacat**cgg** |
| *Nme*Cas9 | *F8* | ggtttctagttgtgacaagaacactggt**gatt** |
| *Mb*Cas12a | *Luciferase* | tcagcagctcgcgctcgttg**taaa** |
| | *RUNX1* | cttggttttcgctccgaagg**taaa** |
| | *VEGFA* | gcccccttcaatattcctag**caaa** |

**Supplementary Table T4: H1, U6 and U6-H1 hybrid promoter sequences**

Sequences are in 5’ to 3’ orientation

| | | |
|---|---|---|
| H1 | Full H1 | catatttgcatgtcgctatgtgttctgggaaatcaccataaacgtgaaatgtctttggatttgggaatcttataagttctgtatgaggaccacggta |
| U6 | Full U6 | gagggcctatttcccatgattccttcatatttgcatatacgatacaaggctgttagagagataattagaattaatttgactgtaaacacaaagatattagtacaaaatacgtgacgtagaaagtaataatttcttgggtagtttgcagttttaaaattatgttttaaaatggactatcatatgcttaccgtaacttgaaagtatttcgatttcttggctttatatatcttgtggaaaggacgaaa |
| miniU6[1] | minimal U6 promoter fused to the 3’ 29 bp of the H1 promoter | gagggcctatttcccatgattccttcatatttgcatatacgatagcttaccgtaacttgaaagtatttcgatttcttggcttataagttctgtatgaggaccacggta |
| miniU6[2] | minimal U6 promoter with its TATA Box replaced by that of the H1 promoter | gagggcctatttcccatgattccttcatatttgcatatacgatagcttaccgtaacttgaaagtatttcgatttcttggcttataagttcttgtggaaaggacgaaa |

**Supplementary Table T5: Primers used for NGS.**

All primers included either the 5' Illumina (forward) or the 3' Illumina (reverse) adapter for NGS sequencing (depicted below in 5'-3' orientation).

| | |
|---|---|
| Illumina 5' Adapter | ACACTCTTTCCCTACACGACGCTCTTCCGATCT |
| Illumina 3' Adapter | GACTGGAGTTCAGACGTGTGCTCTTCCGATCT |

Primer sequences are depicted in 5' to 3' orientation; the 4 nt sequence marked in bold indicate the indices used to multiplex samples for NGS analysis. 5' Illumina adapter sequences are not shown.

| | | |
|---|---|---|
| *EMX1* | fw1 | **AGTC**GGGCCTGAGTCCGAGCAGAAG |
| | fw2 | **AGGA**GGGCCTGAGTCCGAGCAGAAG |
| | fw3 | **GCGA**GGGCCTGAGTCCGAGCAGAAG |
| | fw4 | **CTGA**GGGCCTGAGTCCGAGCAGAAG |
| | rev1 | **GCCG**CAAAAGGGAGATTGGAGACACG |
| | rev2 | **ATCA**CAAAAGGGAGATTGGAGACACG |
| *GRIN2B* | fw1 | **AGTC**AGGACGGCCAACACCAAC |
| | fw2 | **AGGA**AGGACGGCCAACACCAAC |
| | fw3 | **GCGA**AGGACGGCCAACACCAAC |
| | fw4 | **CTGA**AGGACGGCCAACACCAAC |
| | rev1 | **ATCA**GTGTATGCATACTCGCATGGC |
| | rev2 | **GCCG**GTGTATGCATACTCGCATGGC |
| *CCR5* | fw1 | **AGTC**CATTGCTTGGCCAAAAAGAGAG |
| | fw2 | **AGGA**CATTGCTTGGCCAAAAAGAGAG |
| | fw3 | **GCGA**CATTGCTTGGCCAAAAAGAGAG |
| | fw4 | **CTGA**CATTGCTTGGCCAAAAAGAGAG |

| | | |
|---|---|---|
| | rev1 | **ATCA**GAAGGAAAAACAGGTCAGAG |
| | rev2 | **GCCG**GAAGGAAAAACAGGTCAGAG |
| *F8* | fw1 | **AAGG**CCAAGGGTCAGTCTTCTCTATG |
| | fw2 | **AGTC**CCAAGGGTCAGTCTTCTCTATG |
| | fw3 | **AGGA**CCAAGGGTCAGTCTTCTCTATG |
| | fw4 | **GCGA**CCAAGGGTCAGTCTTCTCTATG |
| | rev1 | **ATCA**CATTTATCTGGGAATGGGAGAG |
| | rev2 | **GCCG**CATTTATCTGGGAATGGGAGAG |
| *RUNX1* | fw1 | **AAGG**TACAGGCAAAGCTGAGCAAA |
| | fw2 | **AGTC**TACAGGCAAAGCTGAGCAAA |
| | fw3 | **AGGA**TACAGGCAAAGCTGAGCAAA |
| | fw4 | **CTGA**TACAGGCAAAGCTGAGCAAA |
| | rev1 | **ATCA**CCAGAGGTATCCAGCAGAGG |
| | rev2 | **GCCG**CCAGAGGTATCCAGCAGAGG |
| *VEGFA* | fw1 | **AAGG**TCCAGATGGCACATTGTCAG |
| | fw2 | **AGTC**TCCAGATGGCACATTGTCAG |
| | fw3 | **AGGA**TCCAGATGGCACATTGTCAG |
| | fw4 | **GCGA**TCCAGATGGCACATTGTCAG |
| | rev1 | **ATCA**AGGGAGCAGGAAAGTGAGGT |
| | rev2 | **GCCG**AGGGAGCAGGAAAGTGAGGT |
